# Advancing bronchial tissue engineering: 3D Bioprinting of a Human Bronchial Model Integrating Stromal and Airway Epithelial Compartments

**DOI:** 10.64898/2026.11.27.768368

**Authors:** Albane Carré, Narimène Houmera, Lucie Essayan, Noémie Delorme, Céline Thomann, Alexandra Erny, Gabin Thete, Elise Maurat, Isabelle Dupin, Karen Moreau, Fabienne Archer, Emma Petiot

## Abstract

Advancing *in vitro* bronchial models is essential to overcome the limitations of traditional 2D cultures and animal testing in respiratory research. Most existing systems fail to replicate the complex architecture and stromal–epithelial interactions of native bronchial tissue. In this study, we present an innovative approach employing 3D bioprinting to fabricate a functional human bronchial model that integrates both epithelial and stromal components.

Using micro-extrusion 3D bioprinting, we engineered a stromal compartment composed of normal human lung fibroblasts (NHLFs) and human umbilical vein endothelial cells (HUVECs) within a custom-designed hydrogel. Pre- and post-printing analysis confirmed high cell viability. Extracellular matrix (ECM) synthesis was significantly enhanced by evaluating and selecting a combination of encapsulated cell density, 3D bioprinted construct thickness, and medium change frequency.

Following epithelial seeding with either Calu-3 cell line or primary human airway epithelial cells (hAECs), we demonstrated that the 3D bioprinted stromal compartment supported epithelial attachment, expansion, and maturation into pseudostratified epithelium at the air–liquid interface. Calu-3 3D model functionality was confirmed by a robust mucus secretion while the hAEC 3D model recapitulated key cell types of the native bronchial epithelium, including basal cells, ciliated cells, club cells, and goblet cells, and associated functions (mucus secretion). Qualitative and quantitative methods were used to characterize the composition of epithelial differentiated population on the hAEC 3D and to compare it to the cell population of the native tissue, highlighting its physiological relevance.

This work highlights 3D bioprinting as a powerful technology for generating biomimetic bronchial tissues. The resulting model offers a physiologically relevant and ethically sustainable platform for studying respiratory biology, disease mechanisms, and therapeutic approaches responses, with potential applications in personalized medicine and environmental toxicology.

## Introduction

The bronchial epithelium plays a key role in protecting the airways against inhaled particles and pathogens, principally through mucus-mediated trapping and mucociliary clearance [1]. However, the airways are continuously exposed to infectious and environmental threats [2,3] leading to a wide range of respiratory diseases [4]. Respiratory diseases such as COPD, asthma, cystic fibrosis (CF), lower respiratory infections, and lung cancers consistently span three of the top six global causes of death, accounting for over 7–8 million deaths annually worldwide [5,6].

Despite efforts to improve knowledge on the occurrence, prevention and treatment of bronchial diseases, the development of effective and affordable therapies is impaired by the lack of physio-pathologically relevant bronchial models. Although animal models (mainly rodents) have been used for decades, their physiological differences from human bronchial tissue [7,8] limit predictive accuracy and hinder the successful translation of drug candidates [9]. Furthermore, ethical considerations are driving a shift towards human cell-based, tissue-engineered bronchial models to reduce and replace the use of animal testing in respiratory research [10].

One of the challenges involved in replicating the physiology of the human bronchial epithelium is recreating its biological complexity. The native human bronchial epithelium comprises numerous epithelial cell types with the four main being: basal cells which act as stem-like progenitors responsible for epithelial regeneration; goblet cells which synthesize and secrete mucus to trap inhaled particles and microorganisms; ciliated cells characterised by their apical cilia which facilitate the propulsion of mucus towards the oropharynx for clearance; and club cells which secrete protective and anti-inflammatory proteins involved into immunomodulatory functions and can act as progenitor [1,11]

A range of *in vitro* bronchial models has been developed and deployed to propose alternatives to animal models [12–14]. Nevertheless, each of them still has limitations to fully recapitulate human bronchial physiology and be employed as a relevant model to study bronchial pathologies. Among them, the two-dimensional (2D) monoculture of epithelial cells cultivated at the air-liquid interface (ALI) is one of the well-established bronchial models. Such models rely on culture inserts to grow either Calu-3, a bronchial cell line originating from human bronchial adenocarcinoma, or primary human bronchial (hAECs) or tracheal epithelial cells (hB/TECs) isolated from patients [15–17]. These models have been shown to recapitulate mucus secretion and barrier functions. Moreover, primary cells models can form a pseudostratified structure presenting the diversity of differentiated bronchial epithelial cell types (ciliated, goblet, basal and club cells) while performing the mucociliary clearance function – i.e., mucus secretion and clearance by the cilia – while retaining self-renewal capacity [18]. Other *in vitro* models propose to improve model relevance by implementing 3D organization, as the bronchioïd model presenting a tubular structure [13], and the 3D organoid models [12,19]. Both models demonstrated a 3D organization, complete epithelial differentiation, and the apparition of functions like cilia beating activity and mucus secretion.

While being closer to the native tissue architecture, those models still lack the crosstalk between the bronchial epithelial and its stromal compartments. Indeed, in native tissues, the bronchial epithelium is supported by a thick sub-epithelial compartment of about hundreds of microns composed of the basal membrane and the lamina propria [20,21]. Such a stroma plays a key role in maintaining tissue homeostasis through the activities of both lung fibroblasts and microvascular cells. The synthesis of a complex extracellular matrix (ECM) and its remodeling [22] as well as an endothelial microvasculature ensuring nutrient supply and metabolic wastes removal is essential for tissue function and integrity [23]. The importance of stromal cells in the development, maintenance, and repair of the bronchial epithelium has been increasingly recognized [22,24,25]. Indeed, fibroblast migration towards damage area and secretion of a provisional ECM have been described as early steps in the wound-repair process [26]. In pathologies such as asthma or COPD, lungs fibroblasts have been described to secrete a modified ECM, characterized by increased collagen content and clumped elastin fibers, directly contributing to changes in tissue stiffness [27]. For their part, endothelial cells have been described to promote bronchial epithelium barrier function through biochemical signaling [28]. Thus, relevant *in vitro* models should incorporate the stromal compartment and its interactions with epithelial cells to more faithfully recapitulate the complexity of the native airway tissue.

Aiming to bridge the gap between 2D culture systems and the human *in vivo* tissue, bioprinting technologies enable the biofabrication of tissue models with varied sizes and micro-architectures allowing precise spatial organization, multiple cell types within custom-designed matrices [29–32]. In this study, we applied a micro-extrusion 3D bioprinting strategy to bronchial tissue engineering, building on previous approaches that successfully generated large and complex three-dimensional structures using stromal cells derived from other tissues [33]. Our objective was to develop and optimize a novel biofabrication workflow for constructing a human bronchial model that incorporates both functional stromal and epithelial compartments (Figure 1). The approach involved bioprinting of a stromal 3D compartment composed of primary human lung fibroblasts (NHLFs) and human umbilical vein endothelial cells (HUVECs) embedded in a custom-developed hydrogel blend. Following a maturation phase, epithelial cells were seeded onto the surface of the 3D construct. Two epithelial cell sources were evaluated: a respiratory standard cell line (Calu-3) and primary human airway epithelial cells (hAECs). The study first evaluated the compatibility of the bioprinting process with stromal cell recovery, then characterized stromal maturation through extracellular matrix (ECM) synthesis and remodeling and the apparition of a neo-micro-vascularization. We finally assessed epithelial functionality by analyzing epithelial differentiation and mucus secretion. The study also aims to assess how biofabricated models could complement existing tools available in the scientific community for investigating bronchial physiology and pathology.

**Figure 1:**
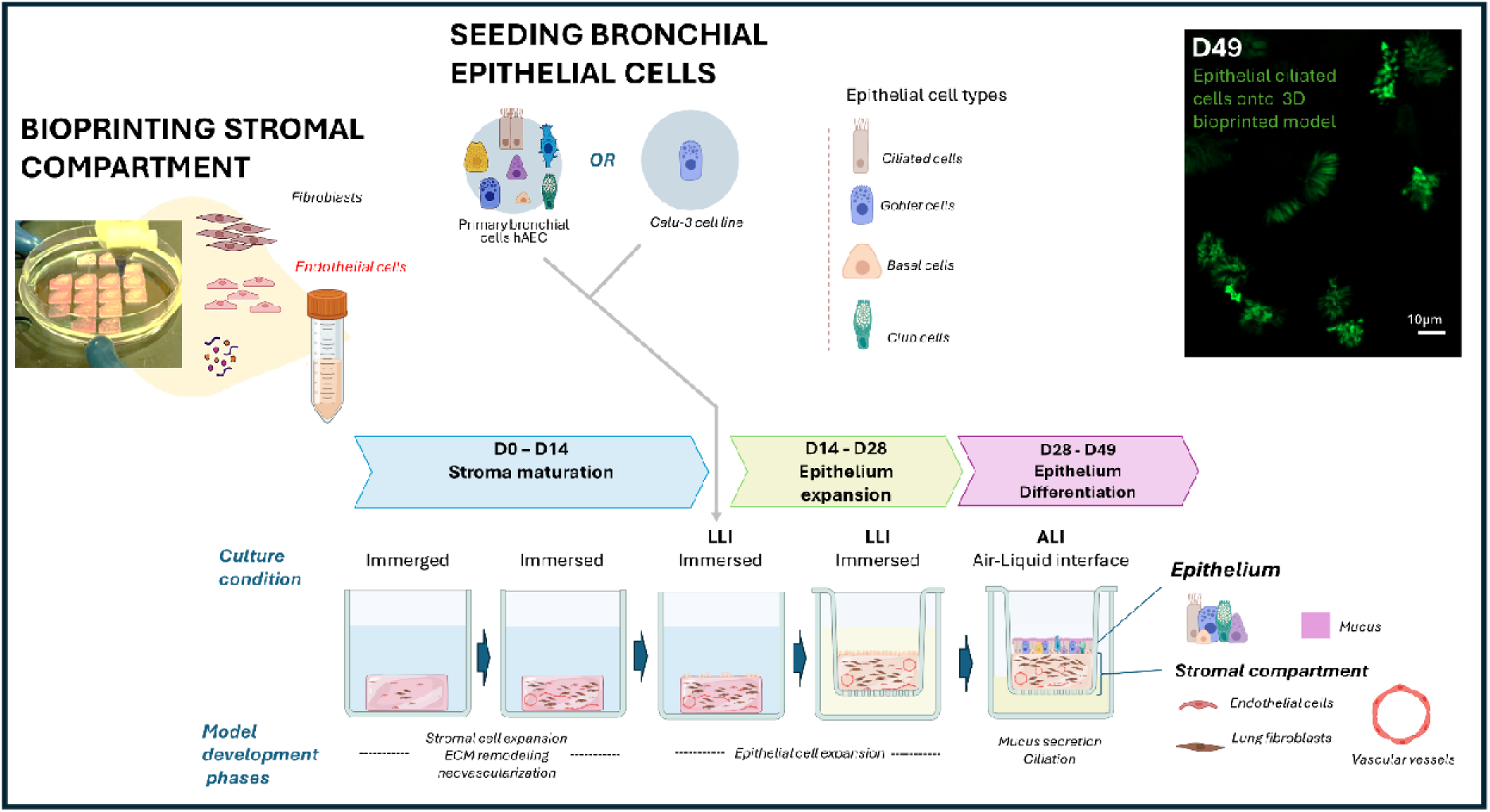
Schematic representation of the workflow used to generate the bronchial models. The stromal compartment was produced by bioprinting a hydrogel containing primary lung fibroblasts and endothelial cells, followed by a 14-day maturation phase under immersed culture conditions (D0–D14), allowing stromal cell expansion, extracellular matrix (ECM) remodeling and neovascularization. Primary human upper airways (bronchial) epithelial cells (hAECs) or Calu-3 cells were then seeded onto the matured stromal compartment. The constructs were maintained under immersed conditions from D14 to D28 to promote epithelial cell expansion, followed by culture at the air–liquid interface (ALI) from D28 to D49 to support epithelial differentiation. At the end of the differentiation phase, the 3D model consisted of a bronchial epithelium supported by a vascularized stromal compartment containing lung fibroblasts and endothelial cells. For models generated using primary bronchial epithelial cells, epithelial differentiation gave rise to the major bronchial epithelial cell types, including ciliated, goblet, basal and club cells.

## Material and methods

### Materials and Methods

#### 1. Cell Culture

Detailed culture conditions for NHLFs, HUVECs, Calu-3, and hAECs, including media compositions, seeding densities, and passages, are provided in Supplementary Material 1. Cells and 3D models were cultured at 37 °C in a humidified 5 % CO atmosphere.

#### 2. Bioink Preparation

The bioink was prepared by dissolving bovine gelatin (G1890, Sigma-Aldrich) (20 % w/v) and very-low-viscosity alginate (A18565, Alfa Aesar) (4 % w/v) in high-glucose DMEM without glutamine and without calcium (entitled DMEM−) supplemented with 10 % FBS, 0.5 % penicillin/streptomycin (15140122, Gibco), and 0.5 % amphotericin B (15290026, Gibco). Fibrinogen from bovine plasma (F8630, Sigma-Aldrich) was dissolved separately at 8 % (w/v) in DMEM−. Both solutions were incubated overnight at 37 °C to ensure complete dissolution. NHLFs and HUVECs were harvested separately, combined at a 1:4 ratio, and resuspended in the fibrinogen solution to achieve a final total cell density of 1.70 × 10 or 2.50 × 10 cells/mL respectively. This cell suspension was then mixed with the gelatin–alginate solution to achieve final concentrations of 2 % fibrinogen, 2 % alginate, and 5 % gelatin (w/v). The cellularized bioink was gently homogenized and loaded into a sterile 10-mL cartridge fitted with a 22-G stainless-steel nozzle (0.41-mm inner diameter, Nordson EFD). Cartridge was equilibrated for 1h at room temperature before printing to stabilize rheological properties.

#### 3. Construct Design, Bioprinting and Stromal Maturation

The 3D stromal compartment was designed as a circular construct with a top reservoir rim and an internal surface area of approximately 1 cm². The bottom thickness ranged from 0.40 to 1.60 mm, while the rim height was maintained at 0.80 mm. Stromal constructs were produced by micro-extrusion in batches of 24 using a BioAssemblyBot® 400 (Advanced Solution). Immediately after printing, constructs were consolidated for 1 h at 37 °C in a solution containing 0.40 % transglutaminase (Activa WM, Ajinomoto), 1 % calcium chloride (T4648, Sigma-Aldrich), and 10 U/mL thrombin (T4648, Sigma-Aldrich). Constructs were then rinsed three times in 0.9% physiological NaCl (Versol solution) and transferred to non-tissue-culture-treated 12-well plates. Stromal maturation was carried out under submerged conditions for 14 days in Mix 2 medium, a 1:1 mixture of FBM and EGM-2+, with medium renewal two to three times a week.

#### 4. Generation and Culture of Epithelialized Bronchial 3D Models

##### 4.1 Epithelial Seeding onto 3D models

After 14 days of stromal maturation, all the culture medium was removed from culture wells and the 3D stromal compartments were incubated for 1 h at 37 °C at 95% humidity to allow partial drying of the internal surface of the rim. Calu-3 cells or hAECs were seeded onto the apical surface at a density of 60,000 cells/cm² in 50µl of their culture medium. The 3D models were incubated for 1 h at 37 °C at 95% humidity to allow epithelial cell adhesion before the addition of culture medium.

##### 4.2 Liquid–Liquid Interface (LLI) Culture and Air–Liquid Interface (ALI) culture of 3D Models

Epithelialized models were cultured at the LLI for 14 days. Calu-3 models were maintained in Mix 3-DMEM/F12, a 1:1:1 mixture of FBM, EGM-2+, and DMEM/F12. hAEC models were maintained in Mix 3-Ex, a 1:1:1 mixture of FBM, EGM-2+, and PneumaCult-Ex Plus, or in PneumaCult-Ex Plus alone. Medium were renewed two to three times a week.

After 14 days at LLI, constructs were transferred to 12-well polyethylene terephthalate inserts and cultured at the ALI for 21 days with 1.5 mL of medium in the basal compartment. Calu-3 models remained in Mix 3-DMEM/F12, while hAEC models were switched to PneumaCult-ALI or Mix 3-ALI, a 1:1:1 mixture of FBM, EGM-2+, and PneumaCult-ALI or Pneumacult-ALI alone. Medium was renewed two to three times a week. The full timeline is provided in Figure 1.

##### 4.3 Epithelial seeding and culture of 2D Epithelial Controls

2D controls for Calu-3 cells and hAECs were seeded on 12-well inserts at 60,000 cells/cm² and maintained at LLI with 0.5 mL of medium in the apical compartment and 1.5 mL in the basal compartment for up to 4 days. Calu-3 2D controls were cultured in Mix 3-DMEM/F12, while hAEC 2D controls were cultured in PneumaCult-Ex Plus until confluence. ALI was then established by removing the apical medium. Calu-3 2D controls were cultured in Mix 3-DMEM/F12 still, and hAEC 2D controls were cultured in either Mix 3-ALI or Pneumacult-ALI alone. The 2D controls were maintained at ALI for 14 days. Medium was renewed two to three times a week.

#### 5. Cell Viability and Epithelial Surface Coverage Assessment

Stromal cell viability in the bioprinted compartment was assessed with calcein-AM and ethidium homodimer-1 of Live/dead viability assay kit (Thermofisher; L3224) and imaged using a fluorescence stereomicroscope (Olympus CKX3) and a fluorescence binocular (Nikon Instrument, ECLIPSE Ts2R-FL). Quantification was performed on day-1 samples using ImageJ, with viability expressed as the number of live cells relative to the total number of detected cells per field of view. Epithelial expansion was evaluated after 1, 7, and 14 days of LLI culture using calcein-AM staining, and surface coverage values were calculated for representative images.

#### 6. Models Histological Processing and Quantification

Three-dimensional constructs were rinsed in PBS1x, fixed overnight at 4 °C in Antigenfix solution (Gauss, P0014), dehydrated through graded ethanol, cleared in methylcyclohexane, and embedded in paraffin. Sections were cut at 5 µm, deparaffinized, and rehydrated. Two-dimensional insert controls were fixed in Antigenfix solution for 1 h at room temperature and processed similarly, with sections cut at 10 µm. The detailed protocol is provided in Supplementary Material 2.

Several labeling were used to characterize the 3D structures. First, the overall tissue architecture was assessed by hematoxylin and eosin staining. Second, formation and overall organization of epithelium was identified by multiplex immunofluorescence using antibodies against CD31 (endothelial cell), collagen I (ECM secretion), and pan-cytokeratin (epithelial cells). Then, detailed observation of deposition of extracellular matrix proteins, including collagen I, collagen III, collagen IV, laminin, was detected by immunohistochemistry. Finally, the identification and localization of epithelial cells sub-populations was assessed by epithelial differentiation markers, such as KRT5, acetylated α-tubulin, CCSP, and MUC5AC, by immunofluorescence. Antibodies, working dilutions, and staining conditions are provided in Supplementary Material 3.

Epithelial thickness quantification was performed manually on H&E-stained sections using ImageJ. Measurements were taken from at least 30 randomly selected points of one sample per condition to ensure representativeness. Immunohistochemistry (IHC) and immunofluorescence (IF) images were quantified using QuPath software and a customed image processing and analysis to automate quantification. Two types of image analysis were developed for IHC and IF treatment. First, cell-counting quantification was performed providing a discrete labelling as for acetylated α-tubulin (α-TUB : ciliated cell counting) and DAPI (nucleus counting in IF). For this process, the QuPath cell-detection algorithm was used. For each marker, cell-detection parameters had to be optimized. Regions of interest (ROIs) encompassing either the epithelium or the supporting extracellular matrix were defined using manual annotation tools at a standardized magnification (100 µm scale). Second, a surface quantification process was performed using pixel classification. This strategy was applied on markers presenting diffuse and spread labelling like IHC ECM markers (collagen I, collagen III, collagen IV, and laminin) and for IF MUC5AC, KRT5, and CCSP markers which prevented a clear cell-by-cell counting. In that case, marker-specific smoothing sigma and threshold parameters had to be settled. All image-analysis parameters are provided in Supplementary Material 4. Quantifications were performed for n=3 to 6 histological slides per condition for IHC ECM markers and n=3 for IF differentiation markers.

#### 8. Three-Dimensional Fluorescence Imaging of 3D Models

For confocal microscopy imaging, the samples were first permeabilized with a solution of 0.2% Triton X-100 in PBS at room temperature for 4 hours. A saturation step (1% BSA and 0.2% Triton X-100 in PBS solution) was then performed at room temperature for 4 hours. Following saturation step, a primary anti-KRT5 antibody is added at 1/500 and incubated for 1 week at 4°C, followed by the rabbit secondary antibody at 1/500 and incubated for 1 week at 4°C. Observations were made using a Zeiss LSM 800 confocal microscope at the CTµ platform. For light-sheet microscopy, the samples were first processed in the same way as those used for confocal microscopy. A clearing method involving lipid removal was then used following the supplier’s protocol (Miltenyi MACS Clearing Kit). This protocol includes firstly, dehydration using successive baths of ethanol of increasing concentration (Eth30% 4 hours, Eth50% 4 hours, Eth70% overnight, Eth90% 4 hours, Eth100% 4 hours et Eth100% overnight) followed by clearing using kit’s solutions for 6 hours. Observation was performed in fresh clearing solution. All these steps are carried out at TA and whilst rotating. Observations were made using a Miltenyi Biotec Blaze light sheet microscope on the PLATIM platform.

#### 9. Mucin Secretion Assay

Apical secretion was collected at day 14 of ALI condition for Calu-3 (n=3) and hAEC (2D n=3, 3D n= 6), and at day 21 for hAEC models (2D n=3, 3D n= 6) by incubating 200 µL of Hank’s Balanced Salt Solution (HBSS, Sigma Aldrich) on the epithelial surface for 15 min at 37 °C. Mucin content was measured by enzyme-linked lectin assay (ELLA) in 96-well plates. Porcine mucin type II was used as a standard, and horseradish-peroxidase-conjugated wheat-germ agglutinin (WGA-HRP) was used as the detection reagent.

A 500-ng/mL mucin standard was serially diluted two-fold to approximately 0.24 ng/mL. Samples were diluted 1:50 or 1:100 in PBS before plating. Plates were incubated overnight at 4 °C, washed three times with PBS containing 1 % gelatin and 0.05 % Tween 20, and blocked with 1 % gelatin in PBS. After washing, plates were incubated with WGA-HRP (0.10 µg/mL) for 1 h at 37 °C. Signal was developed with TMB substrate for 10 min at room temperature in the dark and stopped with 1 M phosphoric acid (10.6 %). Absorbance was read at 450 nm, and PBS blank values were subtracted. The absorbance values are converted to mucin concentrations using the calibration curve. Data are presented as cumulative mucin concentration per surface of epithelium (µg/cm2) ± standard deviation (SD).

#### 10. Transmission Electron Microscopy

Samples were fixed, contrasted, dehydrated, and embedded in resin. Ultrathin sections were imaged using a JEOL JEM-1400 transmission electron microscope at the Ciqle platform. The full protocol is provided in Supplementary Material 5.

#### 11. Flow Cytometric Quantification of Epithelial Populations

3D hAEC samples were first incubated in 55 mM sodium citrate for 5 minutes. Complete dissociation was then achieved by incubation in a solution containing alginate lyase (5 U/mL) and collagenase A (3% in PBS) for approximately 30 minutes at 37°C until full tissue disaggregation. Following dissociation, the entire recovered cell suspension was used for downstream analysis without normalization of cell number between samples. Cells were resuspended in equal volumes of buffer and distributed for staining conditions. A fixable viability dye (Viobility™ 405/520 Fixable Dye, Miltenyi Biotec, ref. 130-130-404) was added at a 1:100 dilution in PBS and incubated for 15 minutes at 4°C in the dark to label non-viable cells prior to fixation. Samples were then washed with FACS buffer (PBS supplemented with 2% FBS and 2 mM EDTA), centrifuged at 2000 rpm for 2 minutes at 4°C, and subjected to fixation and permeabilization using the IntraPrep Permeabilization Reagent kit (Beckman Coulter, ref. A07803). Briefly, cells were fixed with Reagent 1 for 15 minutes at 4°C, washed, and permeabilized with Reagent 2 for 15 minutes at 4°C, followed by an additional wash step.

Intracellular staining was performed in a single-plex format using the following fluorochrome-conjugated antibodies: anti-KRT 5, anti-acetylated α-TUB, and anti-MUC5AC. Corresponding isotype controls conjugated to matching fluorochromes were included for each marker to assess non-specific binding. Antibodies and isotype controls were used at a 1:100 dilution and incubated for 30 minutes at 4°C in the dark. Antibodies and isotype controls references are listed in Supplementary Material 3. Following staining, samples were washed, resuspended, and analyzed by flow cytometry (MACS Quant, Miltenyi Biotech). Data were analyzed using FlowJo software (Tree Star). Standard gating strategies were applied, including exclusion of dead cells based on Viability™ staining, to quantify epithelial subpopulations.

#### 12. Analysis of a Healthy-Lung Single-Cell RNA-Sequencing Reference Dataset

Publicly available single-cell RNA-sequencing data were obtained from the CellHint atlas [34] and accessed through the CZ CELLxGENE Discover platform. The lung compartment of this atlas integrates healthy human lung datasets from multiple studies, including Adams et al. (2020), Travaglini et al. (2020) and Madissoon et al. (2020) [35–38] and provides harmonized cell-type annotations generated with the CellHint framework, which were used directly without further modification.

To establish a physiological reference for the bioprinted lung model, only healthy donor samples were considered. To ensure robust cell-type proportion estimates, the analysis was restricted to donors with the highest numbers of captured cells (range 9000-30000 cells per donor), yielding a final set of n = 10 donors. Global lung cellular composition was first examined across the epithelial, immune, stromal and endothelial compartments. Epithelial cells were then isolated, and the number of cells assigned to each annotated epithelial subtype was extracted to determine their relative abundance. To specifically derive a bronchial epithelial reference, alveolar type I (AT1) and type II (AT2) populations were excluded.

The comparison with the 3D model was restricted to basal, multi-ciliated and goblet cells, the three populations resolved by the cytometry panel. Club cells and ionocytes were detected but excluded from this comparison, as they are not identified by the cytometry markers. Ionocytes were additionally near-absent, with only a few cells per donor. For each cell type, proportions were computed within each donor’s bronchial epithelial compartment and then averaged across donors, and used as a physiological benchmark for positioning the bioprinted 3D hAEC model.

## Results

A sequential biofabrication strategy was used to generate 3D bronchial tissue models by bioprinting a stromal compartment, followed by the seeding of either Calu-3 cells or primary human airway epithelial cells (hAECs) onto a matured stroma. The study first evaluated the compatibility of the bioprinting process with stromal cell recovery. Then, we characterized the stromal maturation through extracellular matrix (ECM) synthesis and remodeling, as well as the development of a neo-microvascular network. Finally, epithelial functionality was assessed by examining Calu-3 maturation and mucus secretion, together with hAEC differentiation, as evidenced by the development of ciliation and mucus secretion. Epithelial differentiation level was characterized for 3D hAEC model and compared to 2D control and to native tissue epithelial population.

### 1. Micro-extrusion biofabrication of the stromal compartment

The biofabrication process employed relies on an in-house developed micro-extrusion bioprinting methodology, previously demonstrated as effective for a range of tissues and cell types [33]. However, a persistent challenge in micro-extrusion bioprinting is maintaining cell viability (i) immediately after printing and (ii) following construct consolidation. During the process, shear stress (a mechanical force induced by hydrogel extrusion through the nozzle) can deform cells, potentially leading to cell death [39,40]

To confirm the suitability of our bioprinting setup for bronchial stromal compartment, we first assessed the impact of wall shear stress (simulated in Supplementary Material 6-Figure 6.1-A & B) on stromal cell viability. Given that the simulated cell recovery rates were significantly lower than those previously observed for other cell types bioprinted in-house [41] , empirical validation of stromal cell viability and recovery was essential. Post-extrusion viability was quantified at 67% ± 4%. This was further complemented by an evaluation of cell viability on Day 1 post-extrusion and consolidation. Live/Dead staining and image analysis revealed a viability of 69± 11%, with cells exhibiting a homogeneous distribution and round morphology (Supplementary Figure 6.1-C). These results confirmed the stability of the viable cell population within the printed stromal construct and validated the feasibility of this biofabrication strategy for producing the bronchial stromal compartment.

### 2. A stromal compartment mimicking the native bronchial lamina propria and basal lamina

To reproduce a stromal compartment capable of supporting epithelial cells and mimicking the native bronchial lamina propria and basal lamina, we first optimized key parameters to ensure vascularization and stromal remodeling. The stromal constructs were cultured for 14 days in a 1:1 blend of fibroblast growth medium (FBM) and endothelial cell growth medium (EGM2+) in agreement with our previous media screening study [42]. The fibroblast:endothelial cell ratio was set at 1:4, based on a preliminary study demonstrating the suitability of this co-culture condition for primary lung fibroblasts and HUVECs. Our goal was to achieve hydrogel degradation and replacement by extracellular matrix (ECM) proteins, such as collagen I, III, IV, and laminin, which are essential for epithelial cell adhesion and proliferation [22,27,43]. We also aimed to observe the development of neo-microvascularization.

Weekly Live/Dead staining revealed a progressive cell densification within the bioprinted constructs, with an observed 100% surface coverage by live cells by Day 7 (Supplementary Figure 6.1-C). Visual assessment suggested that the number and distribution of dead cells remained relatively constant over time, indicating that neither the biomaterial nor the culture conditions induced additional cell death. These results aligned with previous studies using similar biofabrication strategies [33,44,45], confirming the biocompatibility and proliferation-supportive properties of our hydrogel formulation.

To enhance ECM cell synthesis, we screened three parameters: stromal cell concentration in the bioink (1.7 or 2.5 million cells/mL), construct thickness (1.60 mm, 0.80 mm, or 0.40 mm), and medium change frequency (twice or three times a week). These parameters were combined into five experimental conditions (C1–C5), as summarized in Figure 2-A. ECM deposition was qualitatively assessed via H&E staining (Figure 2-B) and quantitatively analyzed using immunohistochemistry (IHC) for collagen I, III, IV, and laminin (Figure 2-C). Image analysis allowed to quantify ECM deposition on tissue slices thanks to surface coverage and to compare the conditions (Figure 2-D). Collagen I, III and IV depositions were quantified for the inner tissue core part corresponding to the native tissue lamina propria, and collagen IV and laminin deposition was quantified for the surface of the construct corresponding to the native tissue basal lamina.

**Figure 2:**
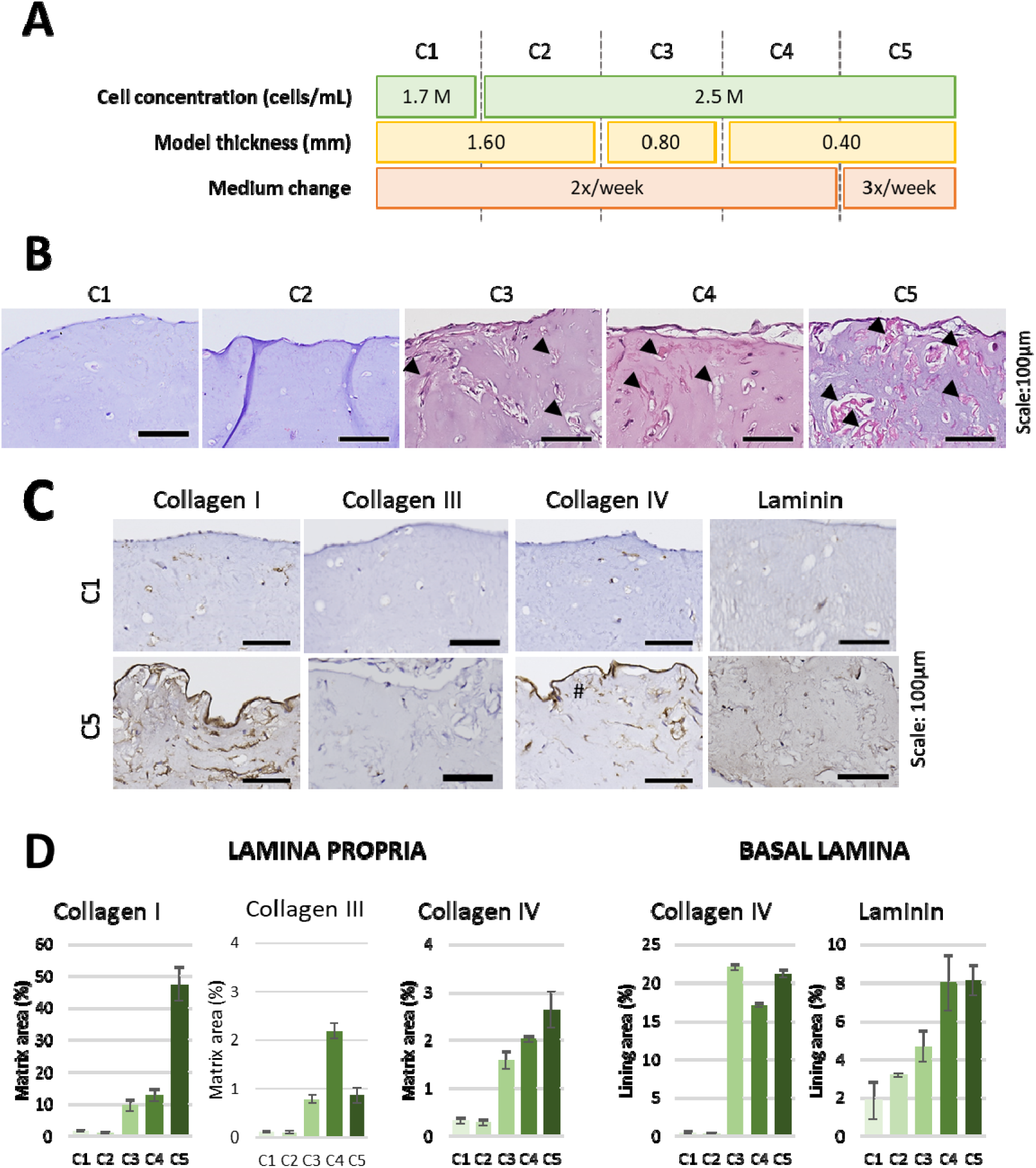
Enhancement of ECM secretion by stromal cells prior to epithelial cell seeding by tuning of the cell concentration, model thickness and medium change frequency. A – representative scheme of the iterative optimisation conditions (C1 to C5). Cell concentration is given for stromal cells, combining NHLF and HUVEC with a 1:4 ratio. Model thickness is the theoretical thickness of the thinner part of the sample, in the centre, as design in the CAD. B – Haematoxylin and Eosin (H&E) staining of each conditions set (C) 14 days after bioprinting. Arrowheads indicate ECM secretion. C – Comparison of collagen I, collagen III, collagen IV and laminin IHC staining for non-optimized protocol (C1) and optimized protocol (C5) at day 14. The collagen IV lining is indicated with #. Scale 100µm. D – Quantification of collagen I, collagen III, collagen IV and laminin secretion for each condition at day 14. Three ROI were defined by tissue slide (n=3) and the fraction of stained area of each ROI was normalized by either the total tissue area for lamina propria, or the surface region manually selected for basal lamina markers.

First, H&E staining showed that increasing the cell concentration alone had no improvement in biomaterial remodeling (conditions C1 and C2). However, reducing the construct thickness from 1.60 mm (C1 & C2) to 0.80 mm (C3) significantly enhanced ECM remodeling, as evidenced by arrowheads on Figure 2-B. Further reducing the thickness to 0.40 mm (C4–C5) did not yield additional improvements.

Thanks to the IHC staining, it was possible to observe additional changes in the sample’s ECM, both for the lamina propria region (inner core) or the basal lamina (surface) regions. First, the low presence of collagen I, III, and IV in the core part of the construct was confirmed for condition C1 (Figure 2-C). As shown by ECM surface coverage quantification (Figure 2-D), the reduction of constructs thickness and the increase of medium renewal frequency from two to three times a week strongly improved Collagen I deposition within the tissue reaching up to ∼50% of the tissue area (30-fold in C5 vs C1). Collagen I is uniformly distributed throughout the construct depth, confirming homogeneous bioink remodeling (Figure 2-C). As for collagen III and IV, the deposition was also improved to a lesser extent in C5 (collagen III: 20-fold vs. C1; Collagen IV: 8-fold vs. C1 in the inner core). Quantification at the surface part of the construct also demonstrated strong improvement of collagen IV and Laminin proteins (respectively 47 and 4-fold vs. C1), thus forming a continuous lining similar to native tissue basal lamina. The presence of these two proteins being essential for further anchorage of epithelial cells, as a consequence, the condition C5—2.50 million cells/mL, 0.40 mm thickness, and three medium changes a week—was selected for subsequent experiments. Overall, construct thickness emerged as the primary determinant of stromal remodeling and ECM deposition, with medium change frequency providing only marginal improvements except for collagen I secretion enhanced by 5 times.

A longitudinal monitoring of ECM deposition for the optimized conditions (C5) demonstrated progressive ECM secretion, with collagen I and IV detectable from day 7 of stromal maturation, while 14 days were necessary to observe a dense and continuous lining of the samples (Supplementary Material 7). Thus, this observation confirmed a necessary pre-maturation step duration of 14 days, prior to epithelial seeding. In parallel, to further investigate the creation of vascularization inside the stromal compartments, an additional experiment was performed using HUVECs carrying the mCherry fluorescent reporter gene. The progressive installation of a vascular network was observed thanks to fluorescence imaging. Full scale model observation was possible thanks to a binocular microscope. After 14 days in culture, the stromal compartment demonstrated neovascular structures of up to 500µm ranges in multiple parts of the bioprinted construct demonstrating the capacity of endothelial cells to self-organize into interconnected structures (Supplementary Material 8.1).

Thus, we demontrated that these culture conditions (2.50 million cells/mL, 0.40 mm thickness, and three medium changes per week, 14 days of culture), resulted in a mature and vascularized stromal compartment presenting ECM lining at its surface. These results allowed us to further develop the model workflow towards epithelialization.

### 3. Epithelial cells anchorage and expansion onto the vascularized stromal compartment

To evaluate bronchial epithelial cell adhesion and expansion onto the bioprinted stromal compartment, Calu-3 cells, or hAECs were seeded onto the surface at a density of 60,000 cells/cm². The models were then maintained at the liquid-liquid interface (LLI) for an additional 14 days to promote epithelial cell expansion and the formation of a continuous epithelial layer.

Epithelial cell coverage was qualitatively monitored using calcein staining. Immediately after seeding, epithelial cells formed discrete patches, distinguishable at high magnification compared to the surface of unseeded stromal compartments (Figure 3-A). To assess expansion dynamics, epithelial cell coverage was quantified from acquired images at different time points. On day 1, both 3D Calu-3 and 3D hAEC models showed adherent epithelial cells with a flat morphology, indicative of early surface colonization, with confluency rates of 11% and 18%, respectively. By day 7, surface coverage increased to 60% for the 3D Calu-3 model and 88% for the 3D hAEC model. Complete epithelial confluency was achieved after 14 days of culture at LLI (corresponding to 28 days of total culture for the stromal compartment; Figure 3).

**Figure 3:**
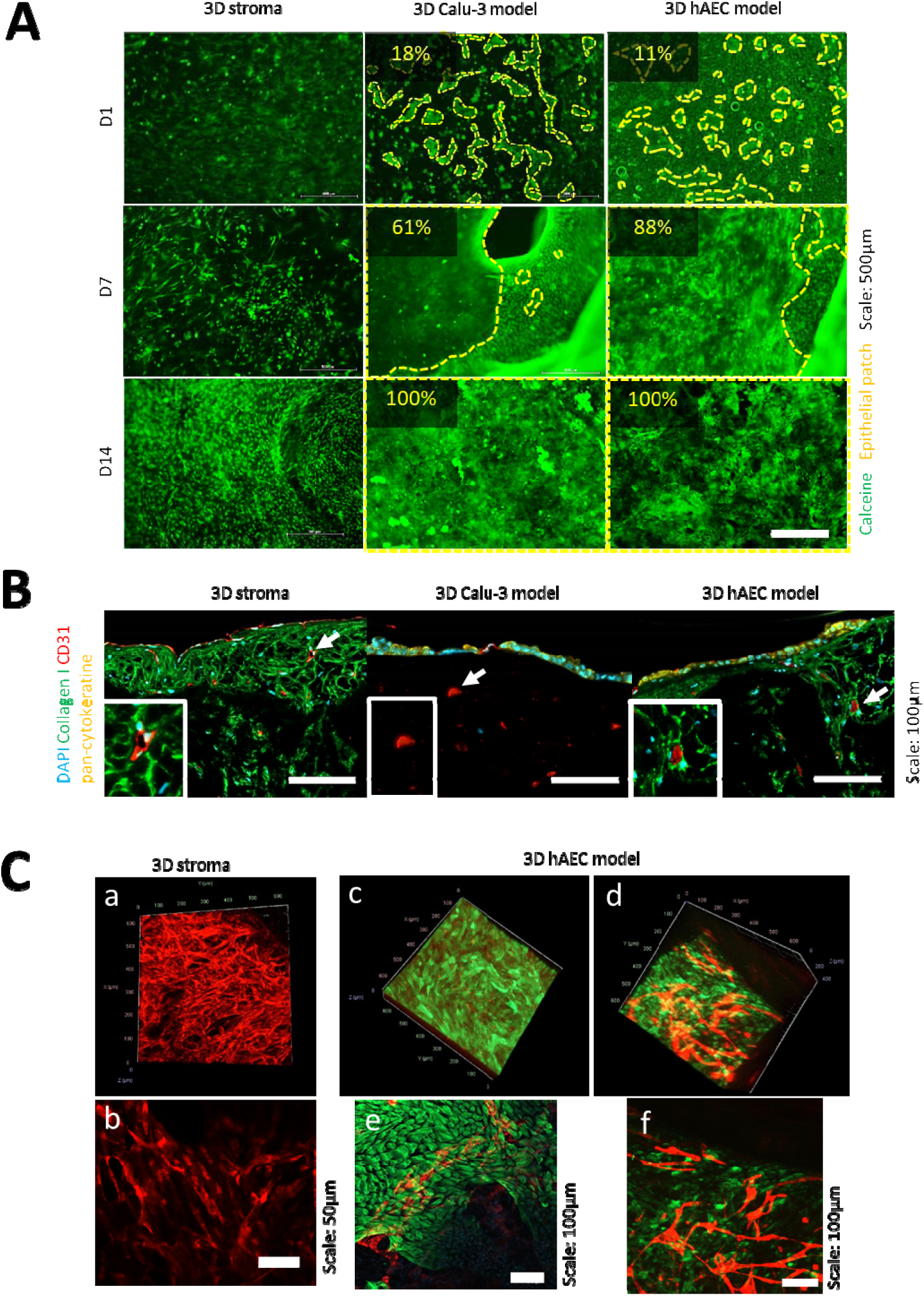
**Epithelial cells expansion and organization onto a vascularized stromal compartment**. A – representative images of live cells development at the surface of the 3D Calu-3 or the 3D hAEC models. The expansion of epithelial patches is highlighted by yellow dotted lines. The epithelial patch surface coverage is indicated on each image. Scale: 500µm. B – Representatives images of the stromal and epithelial cells localization at day 14 of LLI culture, with fibroblasts, endothelial cells and epithelial cells localized by staining of collagen I (green), CD31 (red) and pan cytokeratin (yellow), respectively. Collagen I staining wasn’t performed for the 3D Calu-3 model. All cell nuclei are stained by DAPI (blue). Endothelial structures and lumens are indicated by white arrows and visible in the zoomed image. Scale 100µm. C-Representative images of confocal imaging after 49 days of cultivation for 3D stroma control (a & b) and 3D hAEC model (c,d,e &f). Stromal compartment neo-vascularization identified in red with m-Cherry HUVEC cells, basal epithelial cells identified by KRT5 green labelling. Confocal images represent z-stacks of 45 slices of 4,35µm and a full thickness of 196µm. b,e and f images are orthogonal projections of the 3D volume of samples visualized in 3D on a, c and d respectively.

To confirm the spatial organization of the different cell populations, immunofluorescence staining was performed on histological slices after 14 days of culture at LLI (Figure 3-B). Stromal compartment was characterized with endothelial cells CD31 marker and fibroblast’s collagen I deposition, while pan-cytokeratin staining confirmed the presence of a dense epithelial layer at the surface of both the 3D Calu-3 and 3D hAEC models. In the control non-epithelialized 3D stroma, collagen deposition was observed, consistent with ECM remodeling, and endothelial cells were identified in its core and at its surface. Endothelial cells formed aggregated structures (Figure 3-B, white arrows and magnified images), within the stroma, resembling early lumenization [46]. Notably, a reduced endothelial presence was observed at the surface of the 3D hAEC models compared to the control 3D stroma model (Figure 3-C a&b compared to c&e).

To further investigate the interaction between epithelial and endothelial compartments, HUVECs carrying the mCherry fluorescent reporter gene were tracked in the primary hAEC 3D model. This longitudinal monitoring was run over the LLI and ALI culturing period (Figure 3 C & Supplementary Material 8.2). As shown in Figure 3-C, the fully organized microvascular network established after 14 days of stromal maturation was successfully maintained for 49 days of culture in the control stromal compartment. A progressive increase in vascular network organization (connection numbers and length) was observed.

On the contrary, macroscopic observation of the vascular network in 3D hAEC samples undergoing progressive epithelialization (D15 to D28) and later epithelium differentiation (from D29 to D49) demonstrated a strong interaction between epithelial-microvascular compartment. A gradual reduction of microvascular network vas visible as early as day 21 and prolonged over the 14 days of epithelium growth (see Supplementary Material 8.2). Despite this remodeling process, microvascular structures beneath the epithelium were still visible at the end of the differentiation process (Figure 3-C c-f). Few vessels were positioned right beneath the epithelium, while other structures with branches were visible deeper in the bioprint at approximately 100-300µm depth from the epithelium surface (see Supplementary Material 8.3-B). Vascular network was qualified on five 3D confocal and lightsheet images and quantitative analysis of Z-projected confocal stacks using the Angiogenesis Analyzer plugin in ImageJ. The vascular vessels observed at day 49 were of approximately 7µm diameter and presented lumens (Supplementary Material 8.3-A) . It revealed the formation of branched microvascular structures within the bioprinted construct. These structures are consistent with the characteristic dimensions of human capillaries[47,48] . Measured vascular network on microscopy imaging displayed maximal length of 1380 µm, an average branch length of 21 ± 2 µm, a branch density of 334 ± 63 branches/mm², and a junction density of 624 ± 360 junctions/mm². Interestingly, we should notice that a precise visualization of such a vascular network was only possible through confocal or light-sheet microscopy of the entire bioprinted object and more localized confocal microscopy analyses (see supplementary material 8.4 : video), as histological slices do not permit to observe a representative portion of the tissue in 3D.

Overall, Calu-3 and hAEC cells adhered and expanded on the pre-matured vascularized 3D stromal compartment. To achieve complete functional maturation of those complex models, a culture period at air-liquid interface was mandatory.

### 4. Epithelial compartment differentiation and functionalisation

To thoroughly characterize the differentiation and functional maturation of the primary epithelium in our 3D models, we employed a multi-modal approach combining histology, immunofluorescence (IF), flow cytometry (FC), and electron microscopy (EM). Epithelial maturation was evaluated by H&E staining based on the development of a pseudostratified epithelium and epithelial thickening. Functional maturation was evaluated through mucus secretion in both models. The emergence and organization of differentiated epithelial cell populations, including goblet, club, and ciliated cells, were characterized by immunofluorescence. Cilia formation and tight junction development were assessed by electron microscopy, serving as an indicator of epithelial barrier formation. Finally, flow cytometry provided a complementary quantitative assessment of cell populations distribution. Together, these complementary analyses enabled assessment of epithelial structural organization, cellular composition, and functional maturation, and supported comparison of the 3D models with native human bronchial tissue and other *in vitro* models.

#### 4.1. Bronchial models’ organization

One of the key characteristics of the *in vitro* bronchial models is the development of a pseudo-stratified epithelium, defined as a single layer of cells with nuclei at varying heights, giving the appearance of multiple layers due to differences in nuclear positioning and cell shapes (e.g., columnar, cuboidal, or flask-shaped). Such tissue organization together with epithelial thickening is reported as the first sign of epithelial maturation toward functionality [49,50].

Compared to 2D ALI cultures grown on inserts, both 3D models (Calu-3 and hAECs) reproduced a stratified structure, with greater organization in the case of primary hAEC cells (Figure 4-A and Supplementary Material 9). The thickening of the epithelium, observed *via* H&E staining at day 21 of ALI culture for both models, indicated the transition to a pseudo-stratified epithelium (Figure 4-A). Both models presented an organized structure, with a layered structure resembling native tissue lamina propria, basal lamina and epithelium organization.

**Figure 4:**
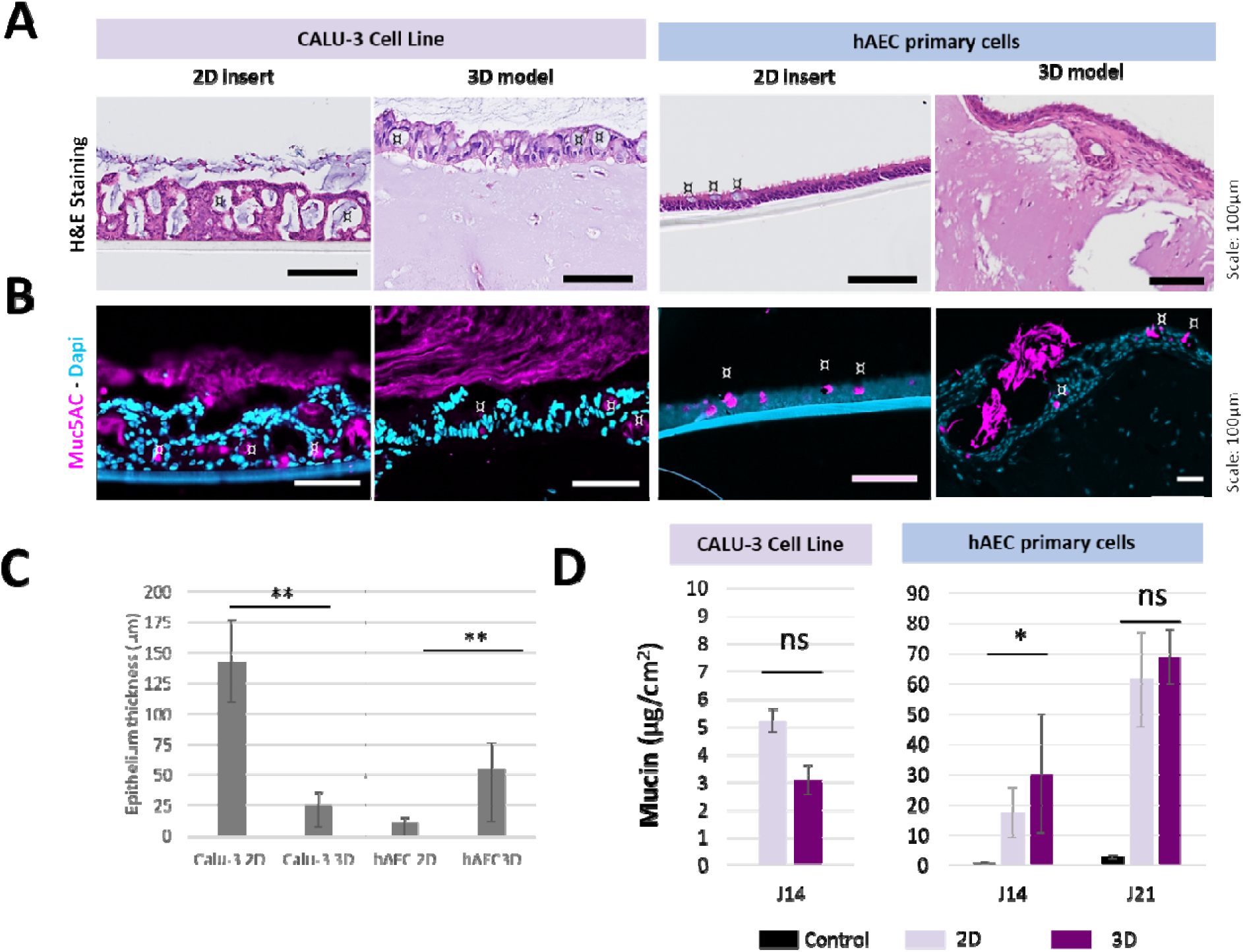
Epithelial cell organization on the 3D models as a pseudostratified epithelium. A – Histology of 3D Calu-3 and hAEC models after 21 days of ALI culture stained with H&E and compared with control 2D inserts. Secretion vacuoles are marked with a ¤. Scale 100 µm. B – Observation of mucus secretion (purple, MUC5AC) by Calu-3 and hAEC seeded in 2D insert or onto the 3D model. All cells’ nuclei are stained with DAPI (blue). Secretion vacuoles are marked with a ¤. Scale 100µm. C – Quantification of model’s epithelium thickness after 21 days of ALI compared to control 2D insert. Measurements were taken from at least 30 randomly selected points of one sample per condition to ensure representativeness. Results are given as mean epithelium thickness in µm ± SD. Statistical analysis was performed using a Kruskal-Wallis one-way ANOVA test; ns: non-significant; **p value < 0.01. D – Mucin secretion quantification in 2D and 3D in both Calu-3 and hAEC models by ELLA assay. Apical secretion was collected at day 14 of ALI condition for Calu-3 (n=3) and hAEC (2D n=3, 3D n= 6), and at day 21 for hAEC models (2D n=3, 3D n= 6). Results are presented as mean ± SD. Statistical analysis was performed using Wilcoxon test, with a p=0.5 between 2D hAEC/3D hAEC at 14 days.

For the 3D Calu-3 model, polarized cuboidal cells of homogeneous height with nuclei at varying levels were observed (Figure 4-A), and no epithelial cell infiltration into the stromal compartment was detected. The Calu-3 3D model epithelial thickness was 63.2 ± 19.9 µm, which was thinner than the Calu-3 2D control inserts (142.5 ± 33.9 µm) (Figure 4-C).

Morphological observation of the 3D hAEC model showed a thick epithelium with homogeneous cell packing positioned onto densely remodeled stroma. Two types of regions could be identified with a basal lamina directly supporting the epithelium and a loser stroma presenting more porosity. Quantification revealed that the epithelia were thicker in 3D hAEC models (52.3 ± 22.9 µm) compared to hAEC 2D controls (40.6 ± 3.3 µm) (Figure 4-C).

#### 4.2. Bronchial models mucus secretion

Regarding functionality, both models were tested for their mucus secretion capacity. Immunofluorescence staining for MUC5AC was performed on models and their 2D ALI controls and attested secretion capacity of the 3D models (Figure 4-B). The associated mucus secretion level was measured at 14-and 21-days post-ALI. For comparison purpose, mucus secretion levels were normalized by epithelialized surface (µg/cm^2^). The assay demonstrated comparable secretion at day 14 in the 3D Calu-3 model (5.23±0.43 µg/cm^2^) to the 2D control (3.08± 0.53 µg/cm^2^) (Figure 4-D).

Similar analysis was performed on epithelial models differentiated from primary hAEC. Mucus secreting cells were also identified through immunofluorescence staining of MUC5AC. Mucus accumulation was visible at different locations onto the bioprint which did not display a homogeneous distribution (Figure 4-B and Supplementary Material 10). The associated mucus secretion levels measured at day 14 and 21 were significantly above negative stroma control performed in the same experiment (Figure 4-D). Mucus was efficiently secreted by the 2D and 3D hAEC models, with an increasing cumulative production over time. If the 2D model (n=3, 30.34±19.78 µg/cm^2^) secreted more mucus at day 14 of ALI compared to the 3D model (n=6, 17.62±8.34 µg/cm^2^) (p=0.5 Wilcoxon test), this slight difference was erased at day 21 of culture (2D hAEC control 69.05±8.97 µg/cm^2^ n=3; 3D hAEC model 61.47±15.59 µg/cm^2^ n=6). Interestingly, the mucus secretion of 2D hAEC insert cultivated in Pneumacult media only, presented mucus secretion values in the range of the 3D stromal compartment while cultivated in mixed medium.

#### 4.3. Bronchial epithelium differentiation level of the 3D hAEC model

First, to characterize the cellular composition of the epithelium in both 2D and 3D models, we applied immunofluorescence staining (Figure 5-A) to confirm the presence of basal cells (KRT5: cytokeratin-5), ciliated cells (α-TUB: acetylated α-tubulin), and club cells (CCSP: Clara Cell Secretory Protein) and to observe the epithelium organization. All cell types were successfully identified. The presence of a confluent layer of basal epithelial cells—previously confirmed by confocal and light sheet microscopy (Figure 3) — was once again verified. The cellular organization observed in the 3D bioprint permit to observe the layered organization of cells, similar to the 2D control epithelium. Differentiation with the 3D bioprinted model favored a higher proportion of either ciliated cells, club cells, or goblet cells depending on the bioprint regions (See Supplementary Material 10). Thus, in the 3D hAEC model, the differentiated cell populations were less homogeneously distributed than in the 2D insert.

**Figure 5:**
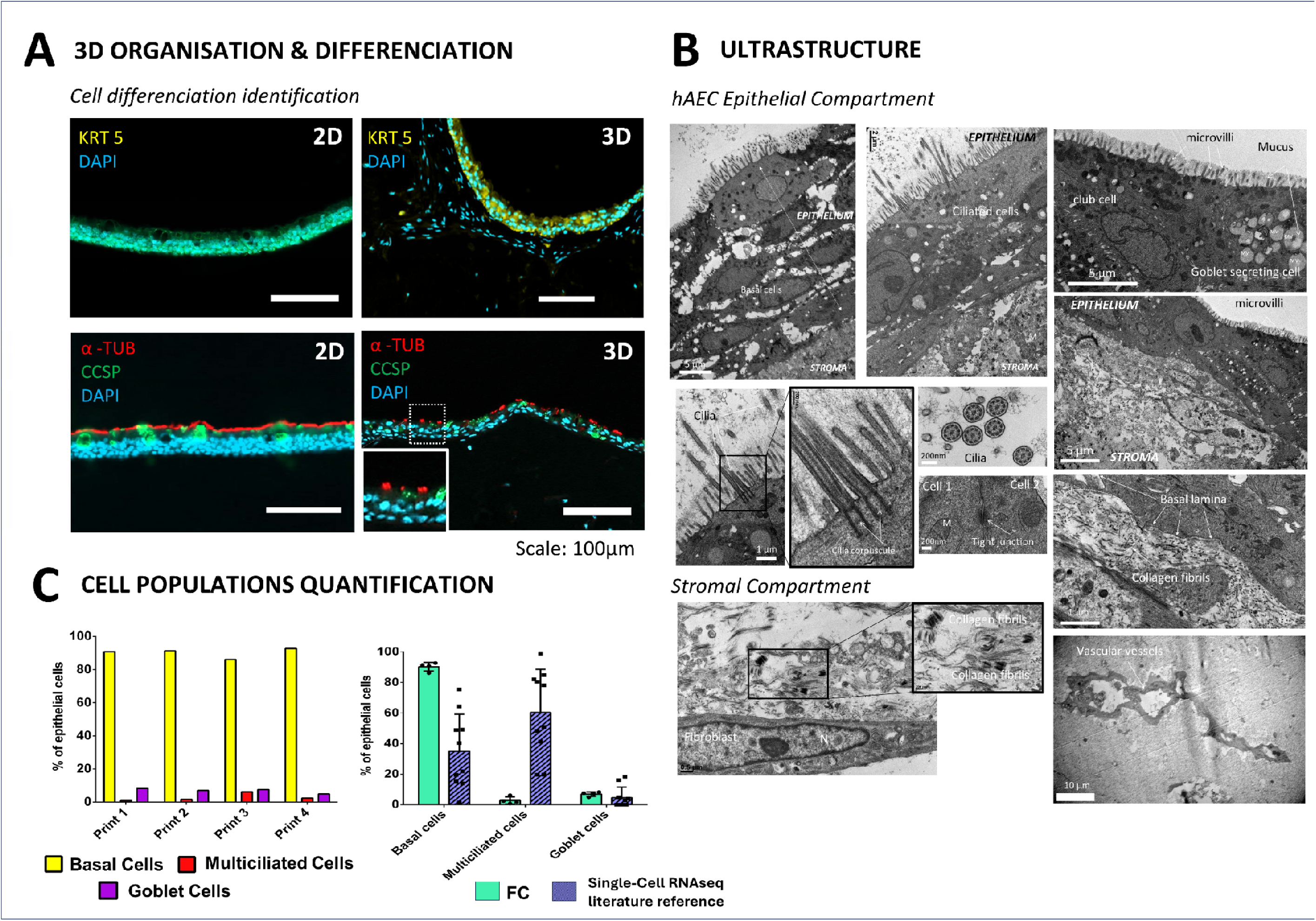
**Description of bronchial epithelium after differentiation of both the 3D hAEC and 2D hAEC control. A-**– Immunofluorescent staining analysis of the presence of differentiated epithelial cells in the 3D hAEC model and the 2D hAEC control. Basal cells (KRT 5+; yellow), ciliated cells (acetylated α-TUB +; red) and club cells (CCPS +; green). All cell nuclei are stained with DAPI (blue). B **- 3D hAEC model ultrastructure at 49 days of cultivation (28 days of ALI) observed by electron microscopy.** Both epithelial and stromal compartment were observed to identify critical cellular ultrastructure’s like cilia and microvilli, basal lamina, collagen fibrils, cell nucleus and endoplasmic reticulum, mitochondria, tight junctions and vascular vessels MV: Mucus vesicles; M : Mitochondria C – Flow cytometry (FC) analysis of epithelial differentiation in 3D hAEC model. Quantification of cells expressing cytokeratine 5 : KRT5+, acetylated α-tubulin+ : α-TUB and Mucine 5AC : MUC5AC+ expressed as percentage of epithelial cells. Data are presented as percentage for individual biological replicates. For each cell population, the mean ± SD of the 4 independent bioprinted 3D hAEC model was compared with literature reference distribution of cell types in the native lung tissue issued from Single cell RNAseq data bases (see Supplementary Material 11)

Ultrastructural analysis revealed features indicative of a polarized and functional tissue (Figure 5-B) on the 3D hAEC model. Ciliated cells were evidenced by the presence of numerous cilia with characteristic axonemal structures. Other cells presented microvilli, signing potential club cells or goblet cells. Mucus secretion vesicles were present within goblet cells, while club cells were observed with their associated secretion vesicles. Observations of several tight junctions and hemidesmosomes confirmed epithelial barrier integrity. Additionally, functional stromal compartment with dense collagen fibrils and organized microvessel ultra structures were identified in the deeper stromal compartment.

Regarding quantification of each differentiated cell populations on the 3D hAEC model, due to the 3D nature of the samples, applying a common quantification based on histological image analysis was complex. This methodology allowed for the quantification of the proportion of ciliated cells, as their discrete labelling (visible bunch of cilia) made it possible to count and normalise by the number of nuclei undeneath. Same methodology applied to club cells and goblet cells did not allow to access to a cell count but rather an epithelial surface coverage, due to the diffuse and spread labelling of CCSP and MUC5AC respectively. Still, by this mean, a mean value of 8.3 +/-4.5 % of ciliated cells was quantified in 3D hAEC model at 21 days of ALI maturation, compared to 36.4 +/-5.2% onto 2D hAEC control inserts. The high percentage variability obtained for the 3D model with such method was related to the heterogeneity in ciliated cell presence depending on the 3D hAEC model regions (see Supplementary material 11).

Consequently, to better assess the cell differentiation level on the full 3D hAEC model, a flow cytometry approach was applied, as implemented previously for other 3D bronchial *in vitro* models [13] (Figure 5-C). This method permitted to access to ciliated cell population to corroborate the above histological quantification, and in addition to quantify goblet and basal cell proportion.

By quantifying cell populations from dissociated 3D hAEC models, we observed a strong predominance of basal cells (KRT5+, 90.2 ± 2.8%), with minor proportions of multiciliated (acetylated α-tubulin+, 2.8 ±2.4%) and goblet cells (MUC5AC+, 7.0±1.5%) within the epithelial compartment. These proportions are consistent with the histological quantification of ciliated cells reported above. Of note, the construct also contains NHLF fibroblasts and HUVEC endothelial cells, whose proportions were not quantified by the cytometry panel. The values above therefore describe the composition of the epithelial fraction, not of the whole construct.

To position the 3D hAEC model epithelial cell composition in a physiological context, we compared it to a published single-cell RNA-seq reference of native human lung (CellHint atlas, [34]). Because whole-lung transcriptomic atlases do not provide a quantification restricted to the bronchial compartment, cell quantification of the bronchial epithelial compartment was extracted from healthy adult donors (n = 10) after exclusion of alveolar population (see Supplementary Material 12). For comparability with the cytometry panel, proportions were computed within each donor’s epithelial compartment, on the same three populations (basal cells, multiciliated cells, goblet cells). In this reference, multiciliated cells were the predominant population (60.4 ± 28.2%, range 20-98%), while basal (35.0 ± 24.2%, range 2-75%) and goblet cells (4.6 ± 6.9%, range 0-18%) showed marked inter-donor variability.

## Discussion

In this study, we established a sequential biofabrication strategy to generate a human bronchial tissue model integrating stromal, endothelial, and epithelial components within a single construct. By optimizing the stromal biofabrication and maturation conditions, we obtained a remodeled extracellular matrix supporting the formation of an organized microvascular network and providing a suitable substrate for subsequent epithelialization. Both Calu-3 cells and primary hAECs successfully adhered, expanded, and formed a continuous epithelium on this bioprinted stromal compartment.

To construct an *in vitro* model, the use of 3D bioprinting introduces specific constraints. Shear stress generated within the nozzle is widely recognised as a major contributor to cell damage and death during the printing process. Assessing this parameter is therefore critical. In our study, we combined *in silico* shear stress modelling with experimental evaluation of cell viability. Our findings showed that, while being used presently on an extensive set of cell types including primary cells [51,52], micro-extrusion bioprinting has a very stringent impact on our primary lung fibroblasts (NHLF) and human umbilical vein endothelial cells (HUVEC). – as they have much lower resistance to shear stress. Still, micro-extrusion was found to be compatible with the stromal cell types we selected, provided that we take into account their inherent differences in resistance to this process and increase the cell concentration.

Achieving ECM synthesis that reproduced the native bronchial tissue composition was proved to be essential to support epithelial health and function [22,24,25]. In healthy tissues, the bronchial lamina propria is characterised by a rich ECM composed of collagen types I and IV, laminin, elastin, fibronectin, and proteoglycans, which collectively provide structural support and regulate stromal–epithelial interactions [53]. These interactions modulate epithelial proliferation, differentiation, and barrier integrity via biochemical and biomechanical cues [54]. Using the bioprinting of fibroblasts and endothelial cells, we manage to achieve a mature stromal compartment capable of strong extracellular matrix (ECM) synthesis with the deposition of major representants of its components. Such capacity could as well be transposed to the modelling of pathologies such as COPD, CF and asthma [27,55] where ECM secretion is deregulated by inducing further enhanced ECM protein deposition.

It is especially challenging to generate such a complex structure of the lamina propria—which includes, in particular, a vascular network—while preserving the long-term stability and macroscopic architecture. Among the parameters tested to enhance ECM synthesis —cell concentration, construct thickness, and media exchange frequency — the thickness of the bioprinted model emerged as the most critical parameter. This highlighted the importance of transport distance of nutrients and oxygen as well as cell waste drainage, to support effective cell growth. It is well established that cell viability, growth, and function within 3D printed constructs are strongly influenced by the diffusion dynamics of the above molecules [56]. Previous studies have already reported growth limitations for fibroblasts [45] and other cell lines [57] beyond millimeters. In line with this, the effective diffusion distance of oxygen *in vivo* is generally limited to approximately 100–200 µm from perfused blood vessels [58]. Our model of 400µm thickness thus permitted to maintain encapsulated cells in a non-limited diffusion range.

Enhanced micro-vascularization is another critical key for the *in vitro* tissues. The microvessel organization and associated ECM remodeling is expected to substantially improve paracrine signaling and promote epithelial functionality [59]. Micro-vascularization structures have already been reported in few engineered lung tissue models [60–62], with structures described having multiple branching points, vessels with lumen the range of 10µm and length of 500 to 1000µm. In our model, the microvascular network generated exhibited a two-layers organization with similar structures as observed by others (branching points up to ∼500 and overall lengths of 450 to 1350µm). We obtained a subepithelial plexus with neo-vessels positioned in the lamina propria and a deeper plexus positioned between 100-300 µm in depth. This structure mirrored the native bronchial architecture as described by Zanini et al.[63] and is also visible for prior studies [64] where the subepithelial plexus is separated from the adventitial plexus by a lamina propria rich in ECM.

Our model seemed to be also particularly suited to study the stromal-epithelial interactions and micro-vascular remodeling. Indeed, one of the most striking observations in our model was the progressive, spatially restricted regression of the microvascular network following epithelial seeding. Such a regression or degradation was observed in the close vicinity to the epithelium, whereas the deeper plexus remained largely preserved. This spatial gradient suggests that the regression is driven by locally acting, likely diffusible epithelial-derived signals, rather than by a uniform loss of endothelial cells. While pre-formed microvascular networks are intrinsically fragile in the absence of perfusion and sustained pro-angiogenic support [65], the localized, distance-dependent pattern observed here points more specifically to an epithelial-to-endothelial cross-talk. Rather than assigning this effect to a defined molecular mechanism, we noted that our system, in which a defined epithelium is placed onto an established vascular network showing a measurable degradation gradient, provides a tractable model to dissect such epithelial-vascular interactions.

The seeding of epithelial cells onto the surface of the bioprinted stromal compartment and their transition to an air-liquid interface (ALI) over a period of 21 days allowed to produce a functional epithelium. The pseudo-stratified epithelium observed in both Calu-3 and hAEC models is a hallmark of mature airway epithelium, characterized by nuclei at varying heights and epithelial thickening. Both 3D models (Calu-3 & hAEC) exhibited efficient mucus secretion comparable to their respective 2D controls. While it is difficult to find quantitative data of *in vitro* mucus secretion in the literature, our results are in agreement with the increase of mucus secretion over time in Calu-3 2D ALI model [66]. This level of secretion was identical in the 3D model, highlighting the added value of 3D models as a screening tool for mucus-related assays such as pathogen adhesion or drug permeability studies [67–70].

Regarding the 3D hAEC model, the primary epithelial cell differentiation was assessed by two complementary approaches. First, histological observation allowed us to confirm the presence of different specific epithelial cell populations and to observe their distribution onto the bioprinted stromal compartment. As expected, epithelial cells were covering the upper surface of the bioprints and ciliated cells, goblets cells and club cells were observed in the upper thickness of the epithelium, while numerous basal cells were forming the basis of the epithelium barrier. The differentiation and structuration of this epithelium was confirmed by ultrastructural analysis (transmission and confocal microscopy) signing a differentiated epithelium, that secretes mucus at is apical side, while being in contact with a mature matrix.

To complement this structural analysis, a total cell analysis to qualify the bioprint’s cell population was applied thanks to flow cytometry. Image-based analysis of histological sections, including manual or automated cell counting, is one of the most accessible and widely used approaches for assessing the proportion of differentiated cells [71,72]. However, in our hands, this approach provided only a limited representation of the overall epithelial cell population ratios. This limitation arose from several sources of variability, including the selection and positioning of regions of interest, as well as operator-dependent interpretation associated with manual or semi-automated analysis. Alternatively, the analysis of the entire bioprint epithelial cell population provided a more comprehensive assessment of the different cell populations [13]. We could quantify the proportion of the ciliated, goblet and basal cells, showing that basal cells were largely represented, which was in accordance with the previous histological observation.

Two non-exclusive features of our system may explain the difference in local differentiation between the 2D and 3D epithelium (low number of ciliated cells and goblet cells). First, the tri-culture medium was designed as a compromise between the requirements of the epithelial, stromal, and endothelial compartments. 3D models were maintained in a 1:1:1 mixture of FBM, EGM-2, and PneumaCult™-ALI to support NHLF, HUVEC and hAEC viability. However, this formulation dilutes the epithelial differentiation medium (Pneumacult^TM^ -ALI) and introduces serum and exogenous VEGF, both of which may interfere or delay the terminal epithelial differentiation. Serum has been associated with a more squamous and less mucociliary phenotype, while reduced HIF1A–VEGFA signaling has been linked to ciliated-cell differentiation, suggesting that sustained VEGF supplementation could limit this process [73]. Thus, although epithelial differentiation was achieved in our 3D hAEC model, the tri-culture medium may favor stromal and vascular maintenance at the expense of complete mucociliary maturation.

Second, the 3D architecture may further limit epithelial differentiation. In conventional Transwell cultures, PneumaCult™-ALI supports differentiation on a thin, porous membrane, allowing direct access to nutrients and differentiation factors from the basal medium. In our model, the epithelium was instead supported by a thick, cell-laden hydrogel. Nutrients and soluble factors must therefore diffuse through the entire stromal compartment before reaching the epithelial layer. This increased diffusion distance may have reduced their availability at the epithelial interface and contributed to the incomplete terminal differentiation observed, in agreement with the diffusion limitations discussed above. This basal-enriched, ciliated-poor composition is a recognized general limitation of *in vitro* 3D airway models, not specific to our system [72,74]. While cell quantification data of the ciliated and goblet cell ratio is difficult to find in literature for 3D models, comparable studies classically demonstrate by histology the creation of a dense epithelial layer presenting discrete ciliated and mucus secretive cells [72,74,75]. On the other hand, studies using primary human bronchial epithelial cells cultured at the air–liquid interface (ALI) on inserts without supporting matrix report that ciliated cells represent approximately 30–70% of the differentiated epithelium, depending on culture medium, donor variability, differentiation time, and quantification method [71,76]. Importantly, the robustness of these estimates varies: Leung et al. [71] quantified ciliated cells by manual counting of apical cells on histological cross-sections from 5–7 donors, whereas Giuranno et al. [76] used α-tubulin IF and counted 500–1,000 cells across five randomly selected fields, providing a more direct cell-marker-based quantification. Nevertheless, both approaches rely on sampled regions rather than whole-insert analysis, and therefore again may not fully capture spatial heterogeneity in ciliation across the different models.

Thus, to position our quantitative assessments, the bioprint model epithelial composition was benchmarked against a healthy human single-cell RNA-sequencing reference for lung (CellHint atlas, [34]). While transcriptomic data do not reflect functional protein expression of bronchial cells, such analysis still permit to classify cell population and access to robust ratios. By this mean we can identify that while the basal compartment is over represented and ciliated compartment under represented compared to the native *in vivo* proportions, the mucus secretion cell compartment is compliant with expected proportions. This observation confirms the value of our model for its mucus secretory phenotype, in line with the efficient quantified secretion of mucus. These features position the model at functional maturation stage that already reproduces stromal and secretory aspects of the native airway. Also, this composition is consistent with an early maturation checkpoint at day 21 of ALI, and whether extended culture would increase the multiciliated fraction remains to be determined. Importantly, the reference itself showed substantial inter-donor variability, indicating that a single native composition does not exist and that this composition should be read as positioning against a physiological range rather than a fixed target. This benchmark provides, to our knowledge, the only quantitative *in vivo* reference available for this model, allowing the maturation gap to be quantified rather than to be left to qualitative appraisal.

Our work highlights 3D bioprinting as a powerful approach for generating biomimetic bronchial tissue models with increased structural and cellular complexity. The 3D Calu-3 model represents a step forward from conventional Calu-3 ALI cultures, which are already widely used for preclinical applications, by incorporating a stromal and endothelial microenvironment while retaining the functional characteristics of the epithelial compartment [77,78]. Beyond this cell line-based model, the integration of primary hAECs provides a complementary and more physiologically representative platform, reproducing the cellular diversity of the healthy human bronchial epithelium. This added values of the 3D hAEC model compared to the 2D model should give more insight into different scientific domains such as i) the study of pathogens interactions with the respiratory epithelium (tropism, cytopathogenicity, replicative cycle, disruption of the epithelia barrier) [79,80], ii) the development of therapeutical approaches by evaluating delivery of muco-penetrating therapeutic or vaccinal vehicles [81], iii) the exposome and the question of the impact of pollutants on the respiratory [82,83]and iv) mimicking pathogenesis involving a matrix remodeling (COPD, Pulmonary fibrosis Asthma,…) [84] .

Finally, unlike organoid-based systems, the models developed here retain a readily accessible apical–basal configuration. This architecture facilitates direct exposure of the epithelial surface and remains compatible with conventional functional assays used to evaluate drug responses, pathogen interactions, or environmental exposures [85]. It may also facilitate the implementation of standardized and potentially higher-throughput experimental workflows. Altogether, this biofabrication strategy provides a versatile human-based platform that combines physiological relevance with experimental accessibility, and potential applications in respiratory disease modelling, preclinical testing, environmental toxicology, and, ultimately, personalized medicine.

## Conclusion

In this study, we developed an advanced 3D bioprinted human bronchial model that integrates, within a single construct, a remodeled extracellular matrix, an organized microvascular network, and a functional airway epithelium. Beyond reproducing epithelial features classically achieved in air–liquid interface cultures, our approach introduces a biologically active stromal microenvironment that supports epithelial attachment, expansion, pseudostratification, and differentiation. The 3D Calu-3 model incorporating a standard bronchial cell line represents a step forward from conventional Calu-3 ALI cultures and could be used for preclinical applications. The 3D hAEC model, incorporating primary bronchial epithelial cells, gave rise to the major bronchial epithelial cell phenotypes, including basal, ciliated, goblet, and club cells, together with mucus secretion and key ultrastructural features of native airway epithelial organization. The incorporation of endothelial and stromal compartments further enabled the emergence of tissue-level epithelial–stromal–vascular interactions that remain poorly represented in conventional airway models.

By combining multicellular complexity, a remodeled tissue-specific microenvironment, microvascular organization, and an experimentally accessible air–liquid interface, this 3D models could be used as *in vitro* platforms to bridge several limitations of conventional epithelial cultures and current 3D airway models. Its modular design also provides the opportunity to independently personalize epithelial, stromal, and endothelial components, offering a versatile framework for investigating bronchial physiology and intercompartmental crosstalk. While further optimization may enhance terminal mucociliary maturation and long-term vascular stability, the present models established a robust foundation for the development of next-generation human bronchial models for respiratory disease modelling, host–pathogen and environmental exposure studies, and therapeutic evaluation.

## Acknowledgments

We gratefully acknowledge the financial support provided by Vaincre la Mucoviscidose, the Gregory Lemarchal Association, and the ANR Shapemed Print@Lung program. We thank Ciqle microscopy imaging platform, specifically Annabelle Bouchardon and Batoule Smatti for their work on triple immunostaining, Bruno Chapuis for his expertise in multiple fluorescence channel imaging and Elisabeth Errazuriz-Cerda for her expertise in TEM imaging. We also thank the PLATIM platform of the SFR Biosciences and the Centre Technologique des Microstructures CTµ” of Lyon 1 Claude Bernard University for access to microscopy equipment and assistance with imaging. We are grateful to Mehdi Maleki (3D.FAB platform) for developing the protocol to evaluate cell survival under shear stress, and to Angélique Champavere (IVPC) for her assistance in developing immunofluorescence staining for the differentiation of primary epithelial cells. We acknowledge the use of Mistral as tool to enhance the syntax and clarity of the manuscript text. The scientific content and interpretations remain the sole responsibility of the authors.

## Author contribution

Conceptualization: AC, KM, FA, EP. Methodology: AC, NH, ID, KM, FA, EP. Investigation: AC, LE, CT, AE, GT, EM, ND, NH, EP. Formal analysis: AC, ND, NH, AE, FA and EP. Writing original draft: AC, LE, NH, ND. Review and editing of the manuscript: all authors. Supervision, funding acquisition and project administration: KM, FA and EP.

---

**Supplementary Material 1:**
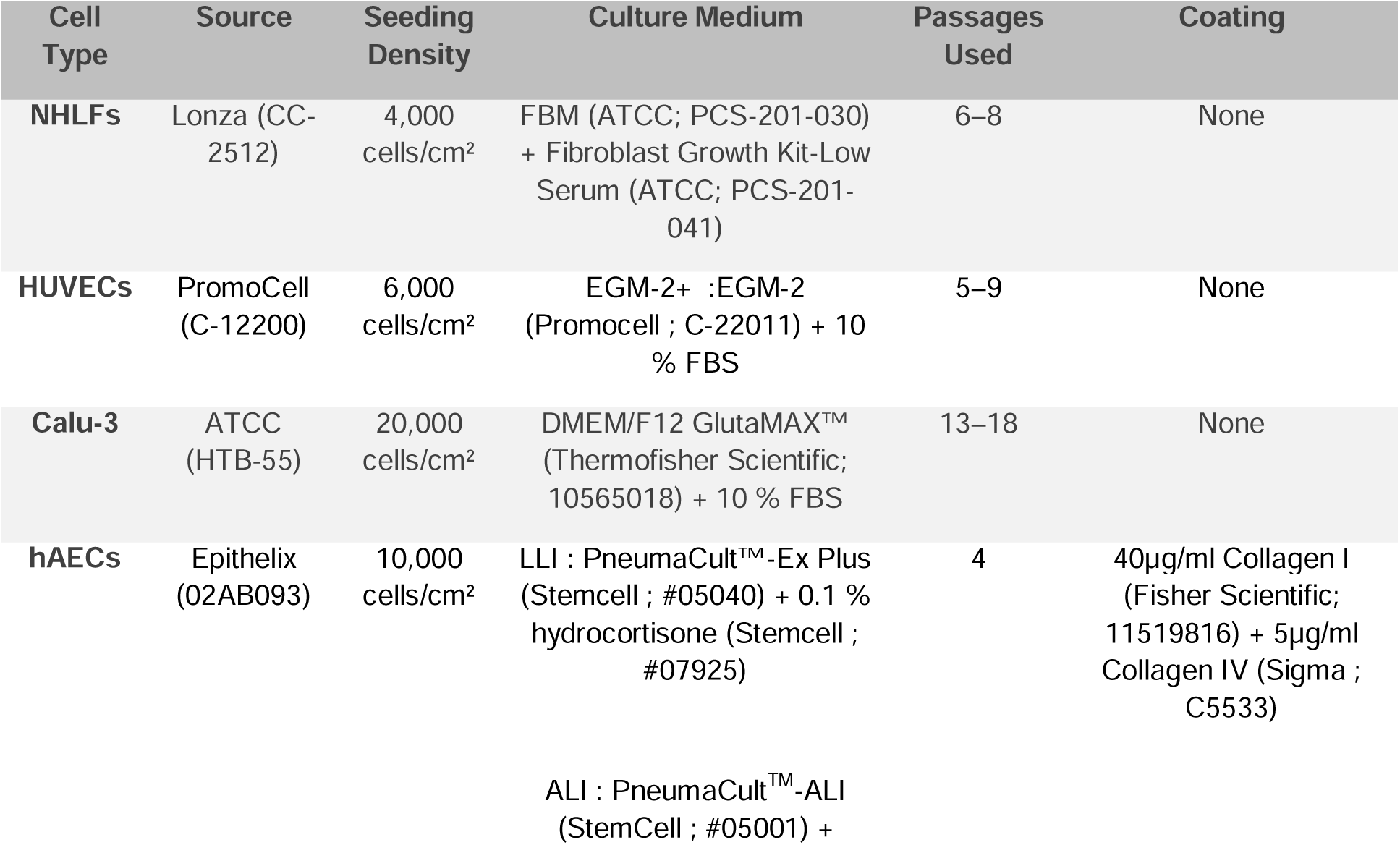
Table describing cell culture conditions.

| **Cell Type** | **Source** | **Seeding Density** | **Culture Medium** | **Passages Used** | **Coating** |
|---|---|---|---|---|---|
| **NHLFs** | Lonza (CC-2512) | 4,000 cells/cm² | FBM (ATCC; PCS-201-030) + Fibroblast Growth Kit-Low Serum (ATCC; PCS-201-041) | 6–8 | None |
| **HUVECs** | PromoCell (C-12200) | 6,000 cells/cm² | EGM-2+ :EGM-2 (Promocell ; C-22011) + 10 % FBS | 5–9 | None |
| **Calu-3** | ATCC (HTB-55) | 20,000 cells/cm² | DMEM/F12 GlutaMAX™ (Thermofisher Scientific; 10565018) + 10 % FBS | 13–18 | None |
| **hAECs** | Epithelix (02AB093) | 10,000 cells/cm² | LLI : PneumaCult™-Ex Plus (Stemcell ; #05040) + 0.1 % hydrocortisone (Stemcell ; #07925)<br>ALI : PneumaCult<sup>TM</sup>-ALI (StemCell ; #05001) + [illegible] | 4 | 40µg/ml Collagen I (Fisher Scientific; 11519816) + 5µg/ml Collagen IV (Sigma ; C5533) |

**Supplementary Material 2:** Table describing Histological Processing Protocol.

| Step | Reagent/Solution | Duration | Temperature | Notes |
|---|---|---|---|---|
| **Fixation for 3D constructs** | Antigenfix (paraformaldehyde-based) | Overnight | 4 °C |  |
| **Fixation for 2D controls** | Antigenfix | 1 h | Room temperature |  |
| **Dehydration** | Graded ethanol (70 %, 90 %, 100 %) | 15 min per step | Ethanol 70 % at room temperature, ethanol 90 % at 35 °C, ethanol 100 % at 55 °C |  |
| **Clearing** | Methylcyclohexane | 25 min | 55 °C |  |
| **Embedding** | Paraffin | 32 min | 60 °C |  |
| **Sectioning (3D)** | - | - | - | 5 µm sections |
| **Sectioning (2D)** | - | - | - | 10 µm sections |
| **Deparaffinisation** | Toluene (2 baths) | 5 min per bath | Room temperature |  |
| **Rehydration** | Graded ethanol (100 %, 90 %, 70 %), distilled water | 5 min per step | Room temperature |  |

**Supplementary Material 3:** Table describing Antibodies for Immunofluorescence, Immunohistochemistry Cytometry, Confocal and lightsheet imaging.

| Target | Antibody | Host Species | Dilution | Supplier | Application | Antigen Retrieval | Blocking reagent | Comments |
|---|---|---|---|---|---|---|---|---|
| CD31 | AF3628 | Goat | 1:100 | R&D Systems | Immunofluorescence | Discovery CC1 Tris-based buffer, 30 min at 95 °C |  |  |
| Collagen I | 20111 | Rabbit | 1:2000 (IHC), 1 :100 (IF) | Novotec | IHC/Immunofluorescence | Hyaluronidase + pepsin | BlockAid (Invitrogen) | Signal development : ImmPACT DAB ; counterstain : Gill’s haematoxylin |
| Pan-Cytokeratin 5 | BM4068 | Mouse | 1:100 | OriGene | Immunofluorescence | Discovery CC1 Tris-based buffer, 30 min at 95 °C |  |  |
| Collagen III | 60315 | Mouse | 1:1000 | Novotec | IHC | Hyaluronidase + pepsin |  | Signal development : ImmPACT DAB ; counterstain : Gill’s haematoxylin |
| Collagen IV | ab214417 | Rabbit | 1:16000 | Abcam | IHC | Hyaluronidase + pepsin | BlockAid (Invitrogen) | Signal development : ImmPACT DAB ; counterstain : Gill’s haematoxylin |
| Laminin | ab78286 | Mouse | 1 :100 | Abcam | IHC | Hyaluronidase + pepsin | 5 % BSA (Euromedex) | Signal development : ImmPACT DAB ; counterstain : Gill’s haematoxylin |
| EN4 | ab8087 | Mouse | 1:10000 | Abcam | IHC | Citrate buffer (pH 6), 20 min at 100 °C | 5 % BSA (Euromedex) | Signal development : ImmPACT DAB ; counterstain : Gill’s haematoxylin |
| KRT5 | CPA1656 | Rabbit | 1:400 | Cohesion Biosciences | Immunofluorescence | Citrate buffer (pH 6), 20 min at 100 °C |  |  |
| Acetylated α-Tubulin | T6793 | Mouse | 1:200 | Sigma-Aldrich | Immunofluorescence | Citrate buffer (pH 6), 20 min at 100 °C |  |  |
| CCSP | 10490-1-AP | Rabbit | 1:200 | Proteintech | Immunofluorescence | Citrate buffer (pH 6), 20 min at 100 °C |  |  |
| MUC5AC | ab24071 | Rabbit | 1:200 | Abcam | Immunofluorescence | Citrate buffer (pH 6), 20 min at 100 °C |  |  |
| KRT5 | CPA1656 | Rabbit | 1:500 | Cohesion Biosciences | Confocal microscopy | Citrate buffer (pH 6), 20 min at 100 °C |  |  |
| Rabbit secondary antibody | ref 11008, Invitrogen |  | 1/500 | Invitrogen | Confocal mycroscopy |  |  |  |
| KRT5 | ab193894 conjugated alexa fluor 488 |  | 1:100 | Abcam | Cytometry |  |  |  |
| Acetylated α-Tubulin | 35652 conjugated Alexa fluor 488 |  | 1:100 | Cell Signaling | Cytometry |  |  |  |
| MUC5AC | NBP2-32732AF750 conjugated Alexa fluor 488 |  | 1:100 | Bio-Techne | Cytometry |  |  |  |
| Isotype control KRT5 | ab199091 (Alexa Fluor 488) |  | 1:100 | Abcam | Cytometry |  |  |  |
| Isotype control Acetylated α-Tubulin | 2975 (Alexa Fluor 488) |  | 1:100 | Cell Signaling | Cytometry |  |  |  |
| Isotype control MUC5AC | NBP1-43319AF750 (Alexa Fluor 750) |  | 1:100 | Bio-Techne | Cytometry |  |  |  |

**Supplementary Material 4 :** Table describing image-analysis parameters.

| **Target** | **Histology method** | **Thresholding method** | **ROI selection** | **Smoothing sigma** | **Threshold** |
|---|---|---|---|---|---|
| **Collagen I** | IHC | 3,3′-diaminobenzidine (DAB) signal | Matrix | 1 | 0.2 |
| **Collagen III** | IHC | 3,3′-diaminobenzidine (DAB) signal | Matrix | 1 | 0.2 |
| **Collagen IV** | IHC | 3,3′-diaminobenzidine (DAB) signal | Matrix | 1 | 0.2 |
| **Collagen IV** | IHC | 3,3′-diaminobenzidine (DAB) signal | Epithelium | 1 | 0.2 |
| **Laminin** | IHC | 3,3′-diaminobenzidine (DAB) signal | Epithelium | 1 | 0.3 |
| **MUC5AC** | IF |  | Epithelium | 1 | 4000 |
| **KRT5** | IF |  | Epithelium | 1 | 4000 |
| **CCSP** | IF |  | Epithelium | 1 | 2000 |
| **Acetylated α-tubulin** | IF |  | Epithelium | 1.5 | 100 |
| **DAPI** | IF |  | Matrix/epithelium | 1.5 | 100 |
| **Haematoxylin** | IHC |  | Matrix/epithelium | 1.5 | 100 |

**Supplementary Material 5: Transmission Electron Microscopy Protocol** Constructs were fixed for 15 min at 4 °C in a 1:1 mixture of culture medium and 4 % glutaraldehyde (EMS-ref 16320), then stored in 2 % glutaraldehyde in 0.2 M sodium cacodylate buffer (EMS ref 11653) (pH 7.4) at 4 °C. Before processing, samples were washed three times for 1 h at 4 °C in cacodylate buffer, contrasted for 1 h at room temperature with Oolong tea extract (EM grade ref GP18-10) in 0.2M cacodylate buffer, and post-fixed for 1 h at room temperature in 1 % aqueous osmium tetroxide (EMS 19192). After three washes in 0.2 M sodium cacodylate and one wash in water, samples were dehydrated through a graded ethanol series (30 %, 50 %, 70 %, 90 %, 100 %) and transferred to propylene oxide (REF). Samples were infiltrated with a resin mixture (Embed 812, EMS , ref 14901 +DDSA, EMS, ref 13701 +NMA, EMS ref 19001) containing 75 % Epon A, 25 % Epon B, and 1.7 % DMP-30 and polymerized for 72 h at 60 °C. Semi-thin sections (1 µm) were stained with methylene blue–Azur II to select regions of interest. Ultrathin sections (∼70 nm) were cut using a Leica UC7 ultramicrotome, mounted on 150-mesh copper grids coated with 1:1000 poly-L-lysine, and contrasted with uranyl acetate and lead citrate. Sections were imaged at 100 kV using a JEOL JEM-1400 transmission electron microscope equipped with an Orius 1000 camera and Digital Micrograph software (version 1.7).

**Supplementary Material 6 - Microextrusion shear stress description and measured effects on stromal cells**

Microextrusion bioprocesses are described as processes inducing partial cell content lysis while applying shear stress on cells as hydrogel flow through the printing nozzle. To describe the starting live cell content in the stromal compartment, description of shear stress distribution and its impact on stromal cell response was characterised.

**Simulation of shear stress distribution during microextrusion printing**

The maximal wall shear stress experienced by the cellularised bioink during microextrusion was estimated *in silico* as previously described (Lemarié et al., 2021). Computational Fluid Dynamics (CFD) simulations were performed using ANSYS FLUENT® 2020 R1 Academic, which applies the Finite Volume Method (FVM). Simulation parameters included a bioink flow rate of 2.35 µL/s, temperature of 21 °C, and nozzle dimensions of 6.35 mm length and 0.41 mm diameter. These data were further processed with the in-house FlowTips® program (Lemarié et al., 2021), which applies to an analytical Poiseuille-based model to calculate shear rate, shear stress, axial and radial velocities along the nozzle. Results are reported as theoretical shear stress (Pa) relative to the distance from the nozzle centre. Fitting of the simulation was visualized with Matlab software.

**Stromal cell recovery yield post-printing**

To measure the stromal cell recovery after microextrusion and describe its detrimental effects the cell content of cellularized bioink, Bioink samples were extruded through bioprinting nozzle and collected in triplicate, weighed (0.4 to 1.6 g per samples) and dissolved at 37°C for 5 minutes after the addition of at least 5 times the weighed sample mass of prewarmed DMEM(-) culture medium. Dissolved solution was centrifuged 5 minutes at 300 g and cell pellet rinsed with 500 µL of warmed DMEM(-). Collected cell suspension was counted on a Malassez hemacytometer with trypan blue to establish both cell viability and cell yields.

**Supplementary Figure 6.1.**
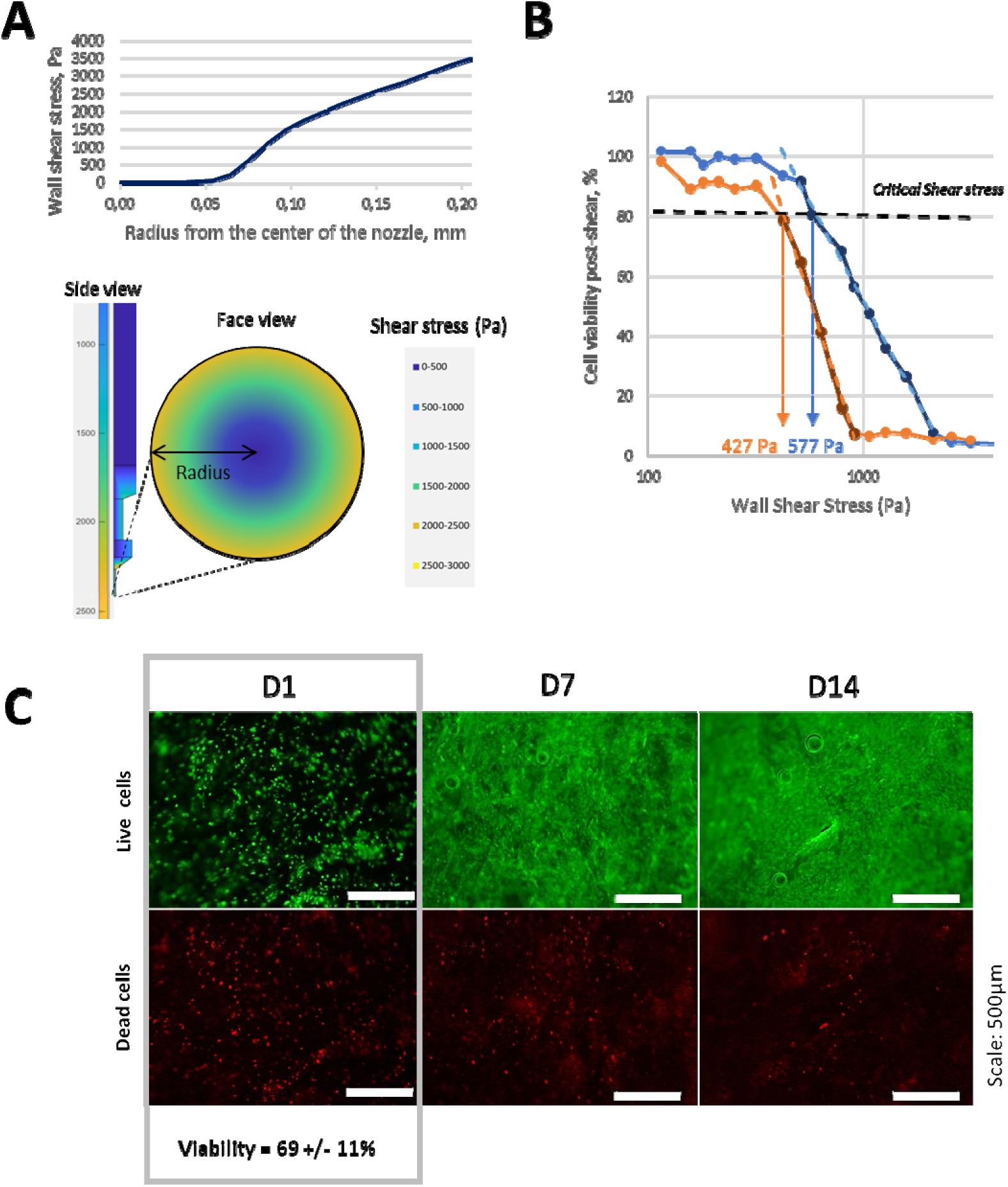
**Micro-extrusion bioprinting effect on stromal cell viability and recovery**. A - Modelling of the distribution of maximal wall shear stress in the bioprinting nozzle expressed in Pascal (Top graph). Modelling parameters: nozzle of 6.35mm length and 0.4mm diameter; bioink flow rate of 2.352µl per sec; bioink temperature 21°C. Schematic representation of the distribution of shear stress experienced by the bioink in the BAB nozzle (Bottom graph). B - Cell viability in response to ranges of wall shear stress applied in a microfluidic device. Critical maximal shear (Critical Shear stress, CSS) was defined as the maximal shear stress cells can sustain retaining 80% of viability. NHLF and HUVEC can sustain 577 Pa and 427 Pa respectively. C – Live and dead cells staining within the bioprinted object at day 1, 7 and 14 after bioprinting. Viability quantification was performed using ImageJ software and expressed as the percentage of live cells relative to the total cell count per field of view. Three independent experiments were conducted, each a minimum of three fields analysed per sample. Scale: 500µm.

**Supplementary Material 7.**
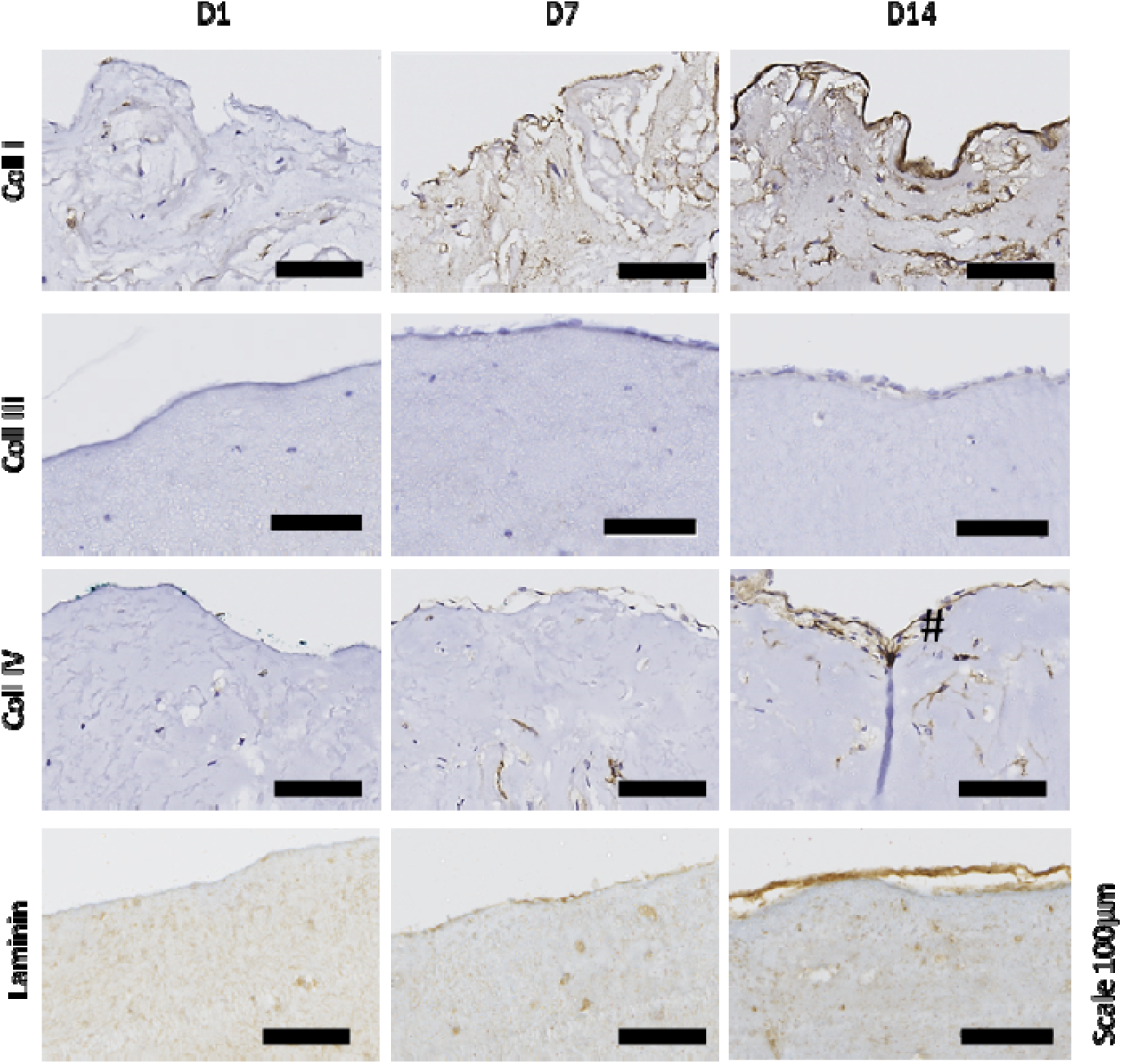
Stromal compartment longitudinal description along its maturation period of 14 days. Representative images of weekly monitored sample’s staining for ECM proteins are presented. Collagen I, Collagen III, Collagen IV and Laminin IHC staining, at day 1, day 7 and day 14 post 3D bioprinting of the stromal compartment. The collagen IV, collagen I and laminin lining are indicated with arrows. Scale 100µm.

**Supplementary Material 8_ 3D bioprinted models microvascularization**

**Supp.8.1.**
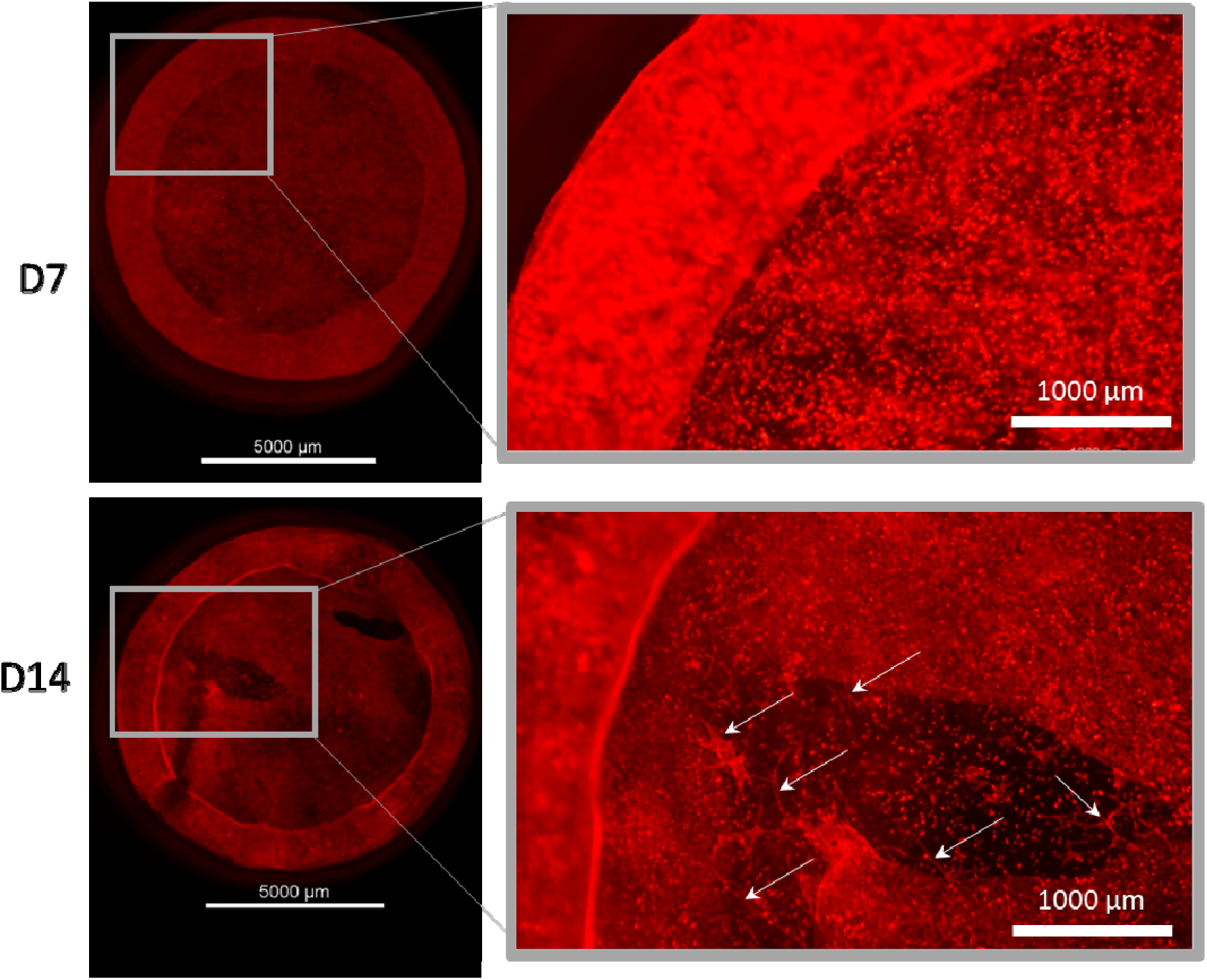
Observation of vascularization apparition in the stromal compartment. HUVEC mCherry cells observed on fluorescence binocular (Stereomicroscope 2CE-SMZ1270, Nikon) after 7 and 14 days of cultivation. Organized vascular structure are observed from D14 and identified as examples by white arrows. Scale 5000 and 1000 µm.

**Supp. Figure 8.2:**
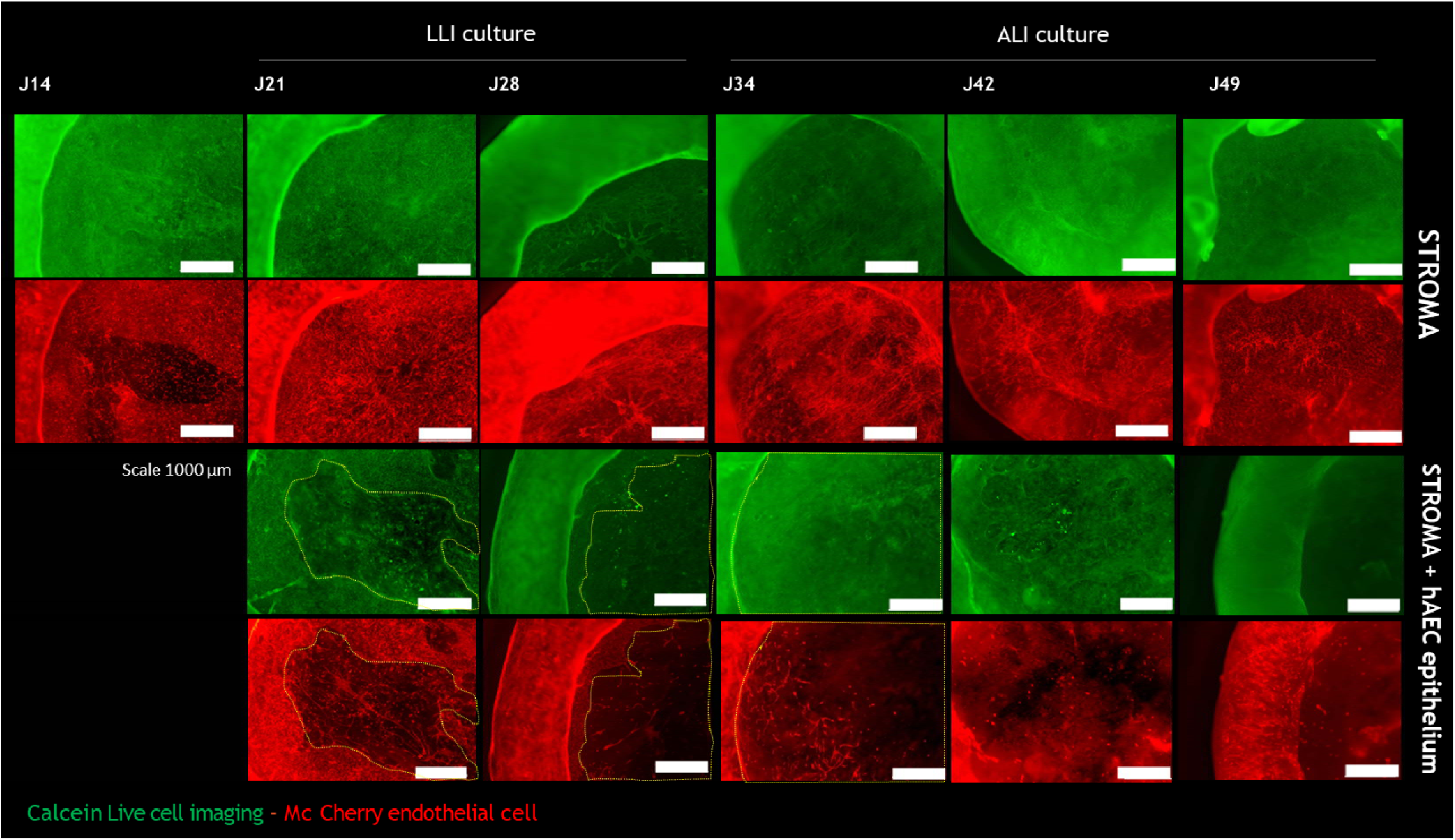
Figure comparing the evolution of neovascularization over the course of growth and differentiation of epithelium compartment from day 14 to day 49. Seeding of hAEC epithelial cell at 15 days and further epithelium installation progressively remodeling / degradation the neighbouring neo organized microvascular network, while stromal compartment without epithelial cells maintains a rich and dense vascular network over the full cultivation period.

**Supp.Figure 8.3:**
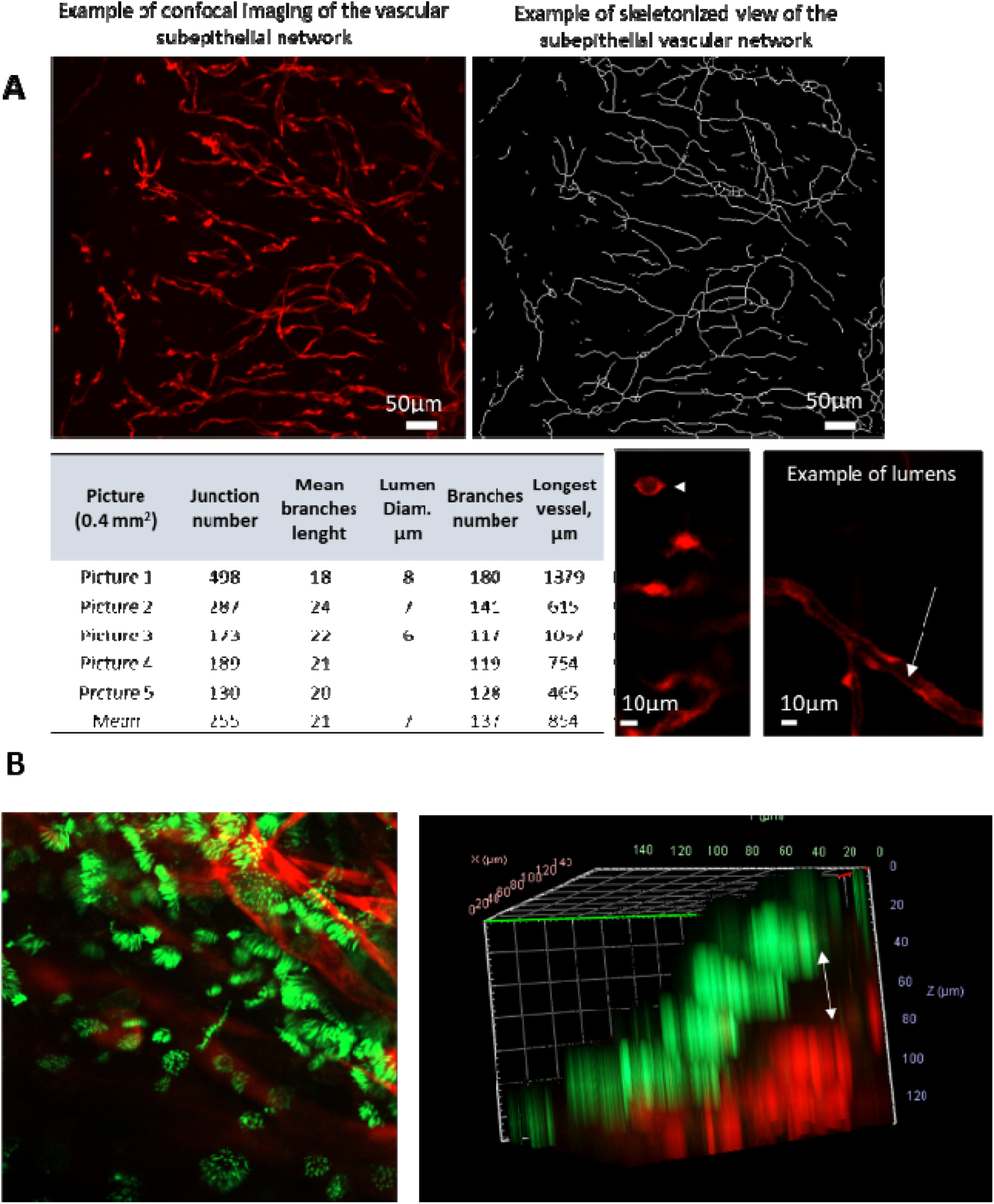
Description of the vascular network characteristics. A. Analysis of vascular network characteristics. These data were obtained using the Skeleton tool in the ImageJ application on images of the objects’ vascular network acquired using a confocal microscope. An orthogonal projection is first carried out, then the images are binarised before applying the Skeleton tool and subsequently the Skeleton Analyse tool. B. Positioning of the subepithelial vascular network. Left : orthogonal projection of the z-stack image of vascularized bioprinted model analysed via lightsheet microscopy; Right : 3D view of the sub-epithelial positioning of the vascular network between 80 to 120µm below the epithelial cells.

**Supp. Video 8.4 : Video of 3D view of bioprinted hAEC model sectors (central part and rim) analysed by confocal microscopy.** Green labelling identifies epithelial cells with cytokeratin-5 epithelial cell marker and red labelling identifies the vascular HUVEC McCherry.

**Supplementary Material 9:**
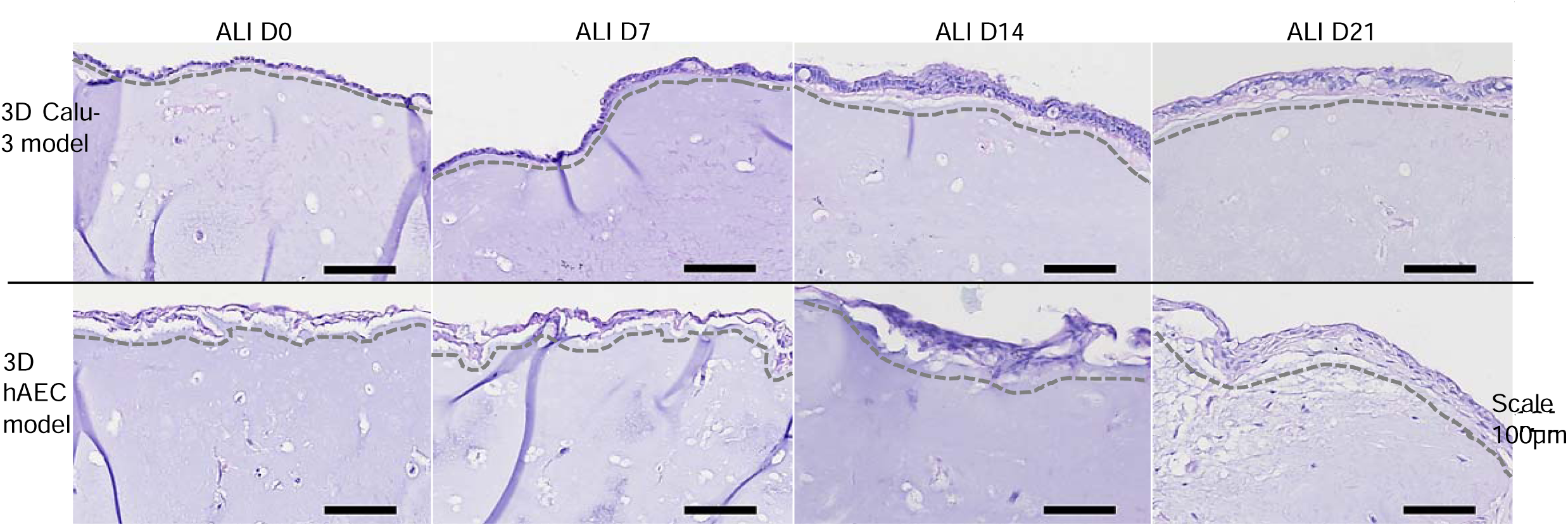
Epithelial cell organization in the 3D models at ALI. Images of the increase of the thickness of epitheliums of 3D Calu-3 model and 3D hAEC model over time. The culture at ALI was conducted up to 21 days. Haematoxylin and eosin staining was performed weekly. The grey dotted line marks the separation between the epithelium and the stromal compartment. (Scale 100µm).

**Supplementary Material 10 :**
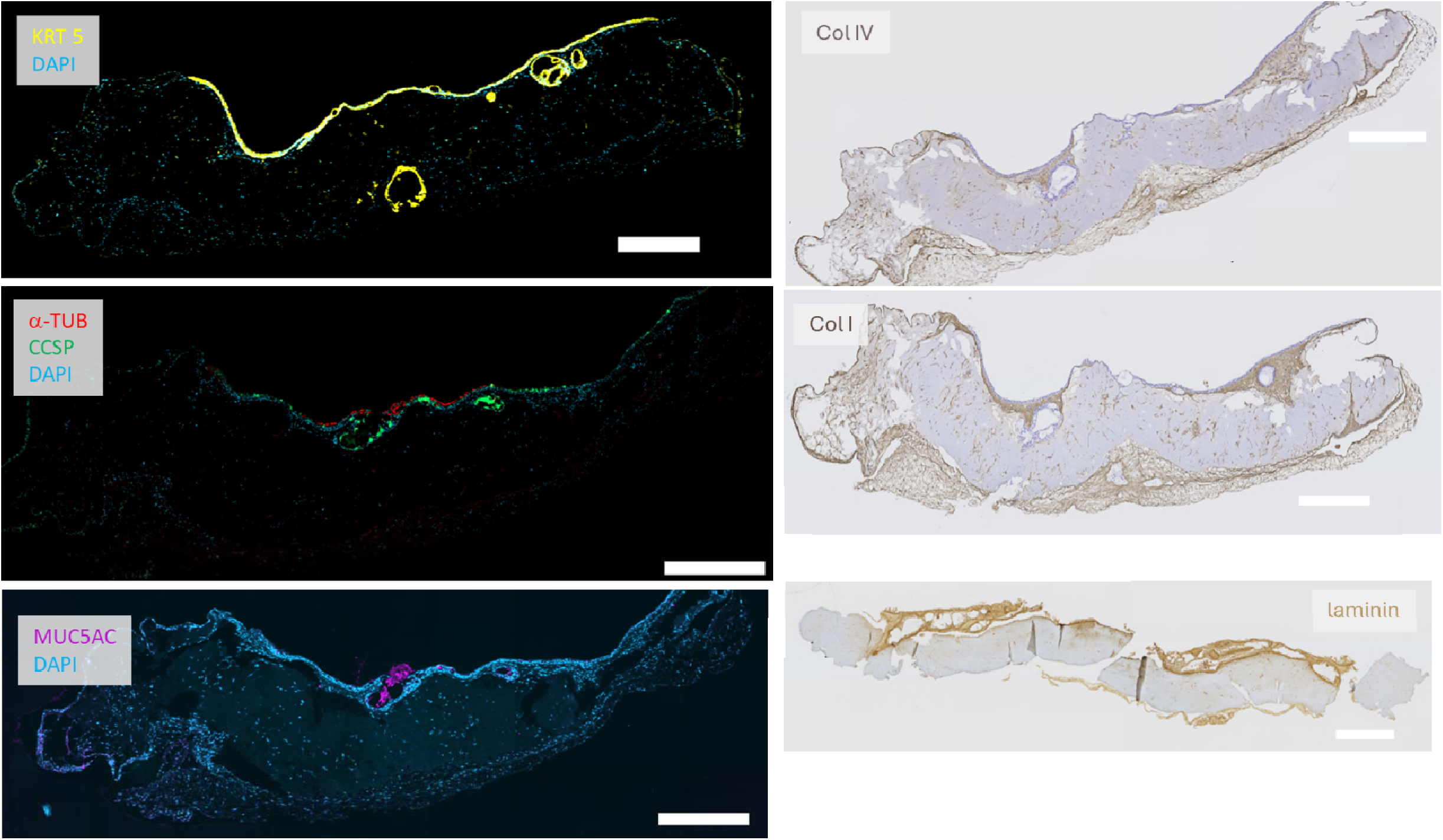
Full view of 3D bioprinted models histological slides for the different labelling. Histological view of 5µm slices of native tissue and 3D hAEC bioprinted model at 21 days of ALI cultivation. Immunofluorescence and Immunohistochemistry staining’s for characteristic labelling i.e. Scale bar represent 500µm.

**Supplementary Material 11 :**
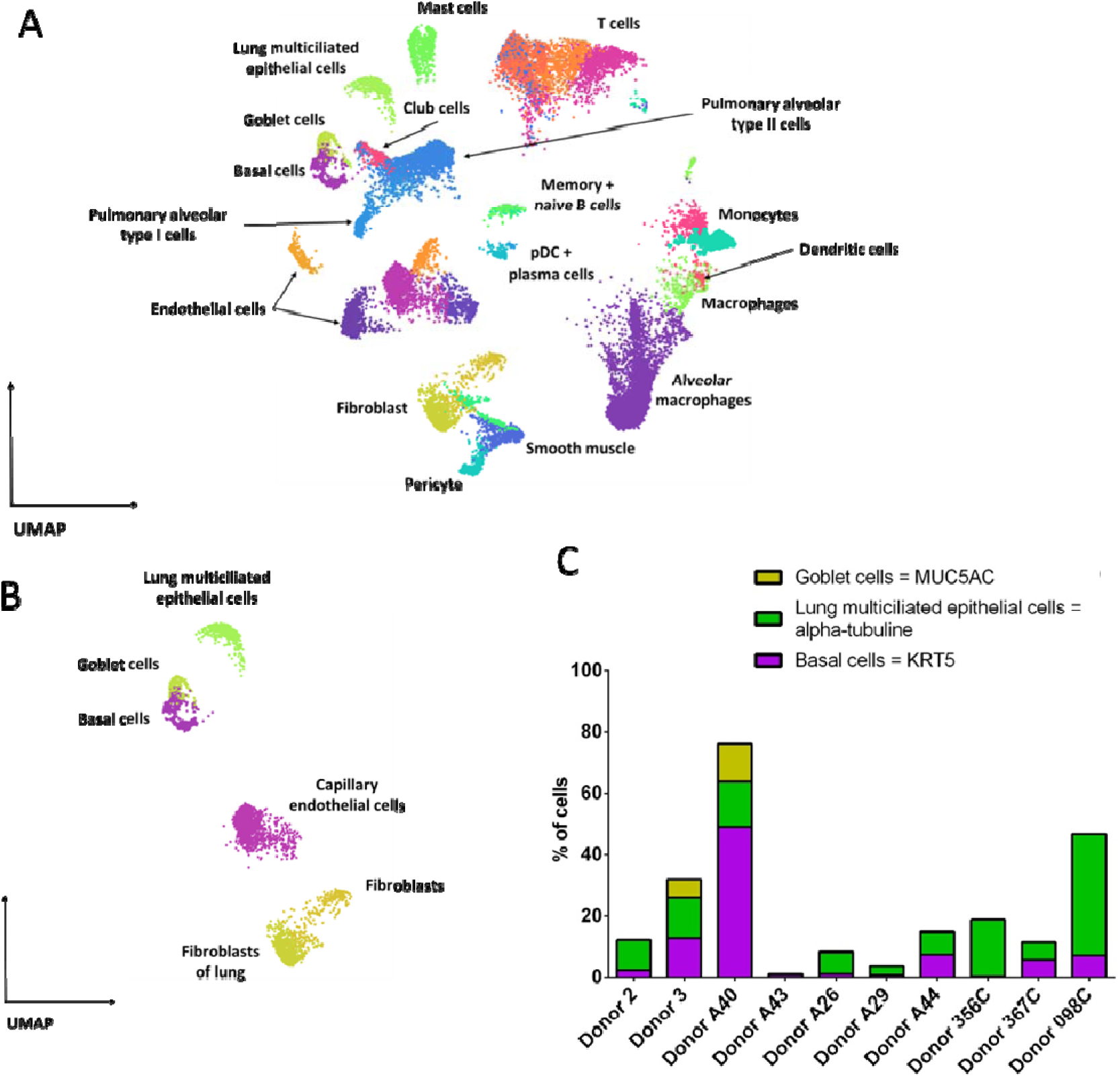
**Data set analysis of the Distribution of cell types in in the native lung tissue (RNAseq data bases)**. A-UMAP representation of a healthy human lung single-cell RNA-seq dataset (donor 3; n=24 655 cells), showing major cellular compartments (epithelial, immune, stromal, endothelial, alveolar). B – UMAP representation of bronchial epithelial cells extracted from a healthy human lung single-cell RNA-seq dataset (donor 3; n=24 cells), highlighting the main epithelial cell populations, including basal cells (n = 563), goblet cells (n = 226), and lung multiciliated epithelial cells (n = 576). C - Comparison of epithelial composition in healthy human lung single-cell RNA-seq reference datasets from different donors, including the three main cell populations (goblet cells, multicillated cells and basal cells).

## Notes

### Competing Interest Statement

The authors have declared no competing interest.

